# Is sediment porosity a confounder in sedimentary DNA records? A test in a temperate semi-enclosed bay

**DOI:** 10.1101/2025.11.19.689188

**Authors:** Tatsuhiko Hoshino, Saki Toyama, Wataru Tanikawa, Masafumi Murayama

## Abstract

Sedimentary environmental DNA (eDNA) has recently been widely used to reconstruct biological communities and to clarify ecosystem dynamics in natural environments. Sediment porosity is thought to influence DNA preservation, but whether it explains variation in sedimentary eDNA beyond depositional setting and burial depth has rarely been tested in natural environments.

We analysed metabarcoding data of 18S rRNA and cytochrome c oxidase subunit I (COI) together with porosity profiles from eight cores (to 6 m depth) collected from the inner and mouth regions of Uranouchi Bay, Japan.

Surface sediments generally showed higher taxonomic richness and read abundance, whereas deeper layers exhibited taxon-dependent decreases in detection frequency. PERMANOVA revealed significant effects of both region and depth on community composition for 18S and COI datasets (p = 0.001).

After accounting for site and burial depth, depth-adjusted porosity explained only a small and non-significant fraction of the variation in community composition (18S: 1.0%, p = 0.74; COI: 4.0%, p = 0.052). In contrast, exploratory class-level analyses showed that diatom classes were negatively associated with depth-adjusted porosity in both markers. Our results indicate that sedimentary eDNA records are primarily structured by depositional setting and burial depth, with porosity contributing little additional explanatory power at the community level within the range observed here. The consistent response of diatoms, however, suggests that porosity effects may be taxon-specific, warranting targeted evaluation in future sedimentary eDNA studies.

**Highlights:**

- Sedimentary eDNA profiles differ between inner– and mouth-bay environments
- Burial depth strongly filters eDNA signals, favouring resistant life stages
- Porosity adds little explanatory power beyond site and burial depth
- Diatom eDNA shows a consistent negative association with porosity

## 1. Introduction

Environmental DNA (eDNA) metabarcoding is widely used as an effective approach for assessing biodiversity in marine environments. This approach enables the non-invasive detection of a broad range of taxa, including species that are difficult to identify morphologically, without the need to collect organisms (Deiner et al., 2017; Pawlowski et al., 2018; Taberlet et al., 2012). Most marine eDNA studies to date have focused on the water column. Sediments, however, have recently attracted attention as archives that integrate ecological information on both present and past biological communities, because DNA accumulates in them over time (Armbrecht et al., 2019). Sedimentary eDNA can complement the microfossil record, as it also captures soft-bodied organisms that lack skeletons and are not preserved as fossils (Lejzerowicz et al., 2013). For example, eDNA recovered from marine sediments has been shown to provide information on changes in marine and terrestrial biodiversity over timescales ranging from glacial–interglacial cycles to millions of years (Armbrecht et al., 2022, 2019; Lejzerowicz et al., 2013).

However, the taxonomic composition of sedimentary eDNA is affected by various processes, including DNA production at the source, transport, deposition, and post-depositional preservation (Nguyen et al., 2024). Hydrodynamic conditions and basin morphology further modify eDNA composition by influencing the balance between terrestrial and marine inputs (Campbell et al., 2025; Herzschuh et al., 2025). Consequently, considerable discrepancies have often been observed between sedimentary eDNA records and microfossil records (De Schepper et al., 2019; Pawłowska et al., 2014).

Although the importance of DNA preservation processes in sediments has been recognised, previous studies on the preservation of sedimentary eDNA have focused mainly on biological and geochemical factors, such as oxygen exposure, organic matter content, and microbial activity (Corinaldesi et al., 2011; Boere et al., 2011; Torti et al., 2018), whereas physical properties such as porosity have rarely been examined. Porosity is a fundamental physical property of sediments that governs solute diffusion and pore-water exchange (Huettel and Gust, 1992). Porosity is therefore expected to affect DNA persistence by regulating the supply of oxidants and nutrients to DNA-degrading microorganisms, as well as the frequency with which DNA encounters microbial cells, extracellular enzymes, and mineral surfaces. Under low-porosity conditions, DNA may also be physically protected through its incorporation into, for example, the interlayers of clay minerals or the interior of microaggregates (Crecchio and Stotzky, 1998; Pietramellara et al., 2009). However, because porosity generally decreases with burial depth, it is difficult to separate the effects of porosity on sedimentary eDNA from those of depth and depositional environment. In addition, porosity varies with other sediment properties, such as grain size, compaction history, and diagenetic alteration, and may therefore affect different taxa in different ways, potentially influencing the interpretation of sedimentary eDNA.

Previous sedimentary eDNA studies have mainly targeted polar, deep-sea, or open continental shelf environments. In contrast, few studies have examined the vertical distribution of sedimentary eDNA over several metres of depth in temperate coastal areas. Including multiple sites that differ in depositional environment and hydrodynamic characteristics is particularly useful for evaluating the relationship between porosity and eDNA composition while accounting for differences in these environmental factors. Uranouchi Bay, Japan, is a semi-enclosed bay in which seawater exchange with the open ocean is restricted by a narrow mouth (Shiraki et al., 1996), and its hydrodynamic characteristics and depositional environments vary from the inner bay to the bay mouth. The bay is therefore a suitable study site for comparing the spatial and vertical distributions of eDNA across different depositional environments and for examining their relationship with porosity.

In this study, we analysed sedimentary eDNA in eight sediment cores collected from the inner bay and the mouth of Uranouchi Bay, a temperate semi-enclosed bay in Japan. For sediment samples from the surface to a depth of 6 m, we performed metabarcoding analyses targeting the 18S rRNA gene and the mitochondrial cytochrome c oxidase subunit I (COI) gene to characterise eukaryotic community composition, and compared sedimentary DNA signals, including the contribution of land plant-derived DNA, between the inner bay and the bay mouth. Furthermore, we quantitatively assessed how much additional variation in eDNA composition could be explained by sediment porosity after accounting for the effects of sampling site and burial depth. We also examined whether specific taxa showed associations with porosity that were consistent between the two genetic markers, to identify candidate taxa for future experimental and geochemical studies.

## 2. Materials and Methods

### 2.1. Study area

Uranouchi Bay is a semi-enclosed tectonic inlet on the Pacific coast of Kochi Prefecture, Japan (33°25′N, 133°25′E). The bay extends approximately 12 km in length (Fig. 1), averages approximately 1 km wide, and is connected to the Pacific Ocean through a narrow mouth. This geomorphology restricts water exchange as confirmed by hydrographic surveys (Shiraki et al., 1996). During the summer stratification, the inflow of oceanic water is largely confined to the mouth of the bay, with limited intrusion into the inner bay. The residence time for half of the inner bay water volume is estimated to be 1–2 months in winter and approximately 2 weeks in summer (Shiraki et al., 1996).

**Fig. 1.**
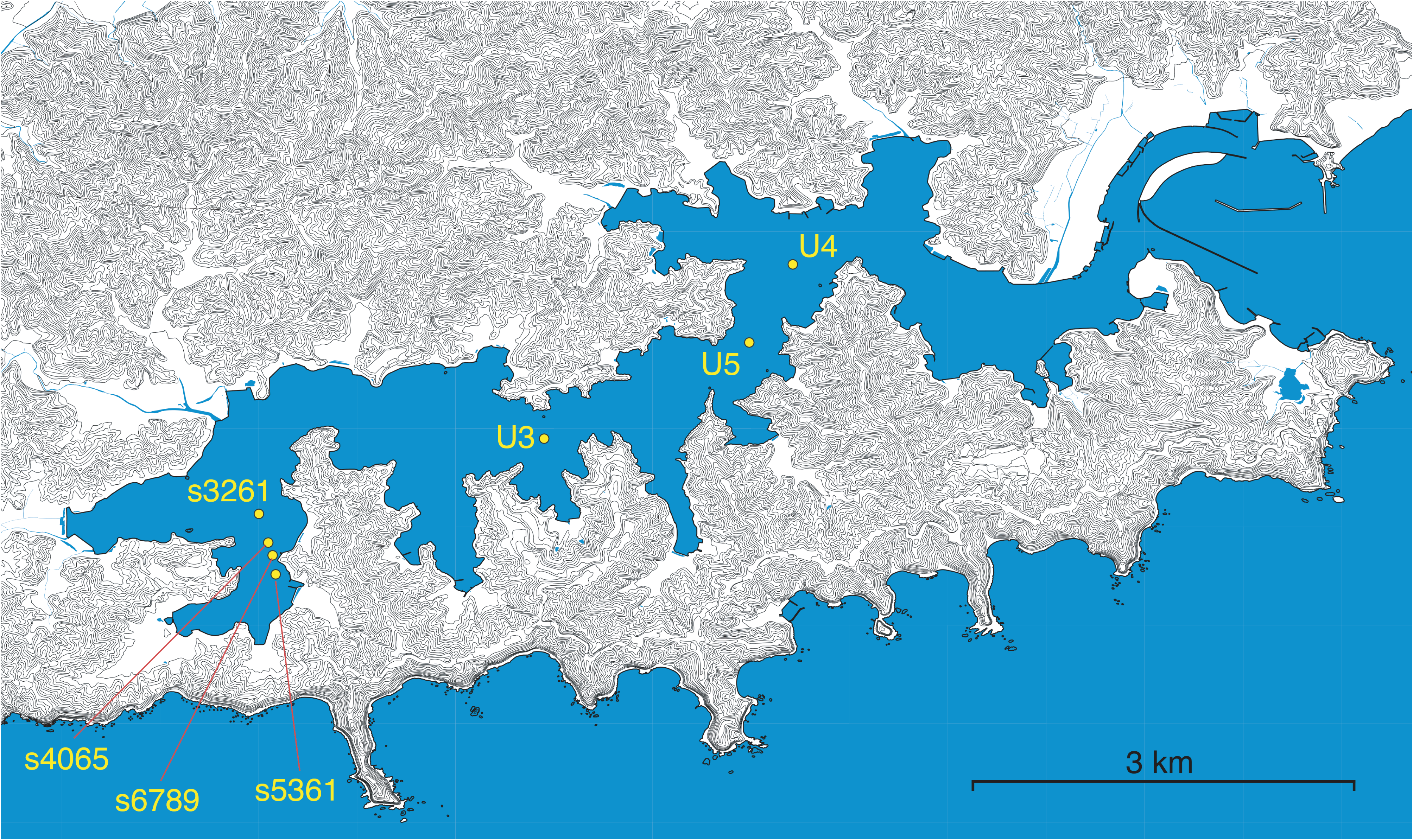
Map of the Uranouchi Bay, Kochi Prefecture, Japan indicating the locations of the sediment core sampling sites. The inner bay sites (UUWs3261, UUWs4065, UUWs5361, and UUWs6789) are located in the central and inner parts of the bay, whereas the bay-mouth sites (U3, U4, and U5) are positioned near the entrance of the bay, close to the Pacific Ocean. This spatial arrangement enabled the comparison of the eukaryotic community composition between the inner bay and bay-mouth environments. The map was created using the QGIS software based on geographic data provided by the Geospatial Information Authority of Japan (GSI).

The catchment surrounding the bay is covered by coniferous (e.g., *Pinus* and *Cryptomeria*) and broadleaf forests, and agricultural land. Water temperature in the bay follows a pronounced seasonal cycle, ranging from approximately 12–14 °C in winter to 28–30 °C in summer; the inner bay exhibits a wider annual range (∼20 °C) than the bay mouth (∼15 °C) owing to its more limited exchange with ocean water. Surface salinity is generally close to oceanic values (∼33–34 PSU) but can drop markedly in the inner bay following heavy rainfall because of freshwater inputs from surrounding rivers (Shiraki et al., 1996). Thermal stratification develops in the inner bay from approximately April to September, leading to hypoxic conditions in bottom waters and enhanced nutrient release from sediments during this period (Shiraki et al., 1996). Aquaculture activities, which began in the 1950s, have further worsened the bay’s water quality and biotic composition through eutrophication (Funaki et al., 2025).

### 2.2. Sampling

Sediment cores were collected across three research cruises aboard the R/V Neptune operated by Kochi University in March 2019 (four sites in the inner bay), March 2021, and June 2022 (three sites near the bay mouth). Seven coring sites were sampled along a gradient from the inner bay to the mouth. Cores of up to 6 m in length were retrieved using a submersible coring system. Immediately after recovery, each core was cut into 1 m sections, either on board or at the wharf, capped at both ends, and stored at 4 °C in the Kochi Core Center repository until subsampling. Subsamples were collected using autoclave-sterilised cut-off syringes and metal spatulas from the cut sections, and ∼5 g of sediment was stored at –20 °C until DNA extraction. A total of 46 sediment subsamples were collected from the eight holes for DNA analysis. Site U3 was cored twice, giving eight holes in total: during the March 2021 cruise, subsamples were collected at 10 cm intervals from 0 to 50 cm depth, and during the June 2022 cruise, subsamples were collected at 100 cm intervals from 100 to 600 cm depth. Coring to 6 m captured a range of post-depositional conditions, from recent surface sediments to older, compacted layers. We did not establish an age–depth model or reconstruct temporal changes in community composition; instead, burial depth was treated as a physical parameter reflecting post-depositional conditions.

### 2.3. Porosity measurement

Porosity was measured using two approaches selected according to the sediment type and core architecture of each site. For the inner-bay cores (UUWs3261, UUWs4065, UUWs5361, and UUWs6789), which consist of fine-grained, cohesive sediments amenable to discrete destructive subsampling, porosity was measured directly on samples collected at 5 cm intervals following the procedure described in Blum (1997). Porosity was calculated from the pore volume, determined from wet and dry weights measured with an electronic balance, and the grain volume measured using a commercial helium gas pycnometer (Pentapyc 5200e; Quantachrome Instruments, FL, USA). The uncertainty of the pycnometer-derived porosity, estimated by propagating the standard deviation of five repeated grain-volume measurements, was 0.03–0.05 vol%.

For the bay-mouth cores (U3, U4, and U5), which were retained as whole-round cores, porosity was estimated non-destructively from X-ray computed tomography (CT) image data (Aquilion PRIME Focus Edition, Canon Medical Systems Corporation, Japan) using a regression model relating CT number to porosity following the method of Gupta et al. (2019). This calibration was established from inner-bay cores where porosity had been independently measured using gravimetric and pycnometric methods. CT-derived porosity was averaged over 10 mm depth intervals using 0.5-mm-spaced CT slice data, and its uncertainty was estimated as the standard error of a 21-point moving average, yielding an absolute uncertainty of 0.1–0.5 vol%. Although pycnometer-derived porosity has higher analytical precision, CT-derived porosity was used for comparison with eDNA community composition because it provides nearly continuous depth profiles at depths corresponding to the eDNA samples.

### 2.4. DNA extraction, amplification, and sequencing

DNA was extracted from each sediment sample using the DNeasy PowerMax Soil Kit (QIAGEN, Germany) following the manufacturer’s instructions with minor modifications, as described previously (Hoshino and Inagaki, 2019, 2024). The average DNA yield was 7.2 ng/g-sediment. As the DNeasy PowerMax Soil Kit employs mechanical and chemical cell lysis, the resulting DNA extract comprises both intracellular and extracellular fractions, consistent with standard practice in sedimentary eDNA studies (Capo et al., 2021).

Two complementary genetic markers were selected to maximise taxonomic coverage of eukaryotic diversity: the V9 region of the 18S rRNA gene provides broad phylogenetic coverage across all eukaryotic lineages, including protists, fungi, and metazoans, whereas the COI gene offers higher taxonomic resolution, particularly for metazoans. Both markers were amplified: (i) the V9 region of the 18S rRNA gene using primers Euk1391f/EukBr (Stoeck et al., 2010), and (ii) a fragment of the mitochondrial cytochrome c oxidase subunit I (COI) gene using primers COX-IntF/COX-dgHCO2198 (Leray et al., 2013; Meyer et al., 2005).

PCR amplification was performed using MightyAmp DNA Polymerase (Takara Bio, Shiga, Japan). PCR negative controls were included in amplification runs, and no visible amplification products were detected in any of the negative controls upon agarose gel electrophoresis. Amplicons were purified using NucleoMag magnetic beads (Takara Bio, Japan), quantified, pooled at equimolar concentrations, and sequenced on an Illumina MiSeq platform using a 2 × 300 bp run (MiSeq Reagent Kit v3).

### 2.5. DNA data analysis

Sequences were processed separately for the 18S and COI datasets. Demultiplexed paired-end reads were merged using USEARCH v11 (64-bit version; www.drive5.com/usearch/). Reads were quality filtered (maxEE ≤ 1.0), and sequences containing ambiguous bases (N) were removed. Quality-filtered reads were dereplicated and denoised into zero-radius operational taxonomic units (zOTUs) (Edgar, 2016).

Taxonomic assignments of 18S sequences were performed in Mothur v1.48 (Schloss et al., 2009) against the SILVA 138 SSU database (Quast et al., 2012) with a bootstrap support threshold of 80%. The COI sequences were assigned against the MIDORI2 GB248 database (Leray et al., 2022). Group-level differences in the relative abundances of major taxa (phylum level) between the inner– and mouth-bay sites were tested using the Mann–Whitney U test.

For β-diversity analyses, libraries with fewer than 5,000 reads were excluded, a threshold chosen to balance sequencing depth against sample retention (Fig. S1). The remaining samples were rarefied to 5,000 reads to standardise the sequencing depth, and read counts were subsequently Hellinger-transformed (square root of relative abundance). Bray–Curtis dissimilarities were then computed, and community patterns were visualised using principal coordinate analysis (PCoA) in two dimensions, with the axes annotated by the percentage of variation explained. To test the significance of community differences between regions (inner bay vs. mouth) and depth categories (shallow: ≤50 cm; deep: >50 cm), permutational multivariate analysis of variance (PERMANOVA) was performed on Bray–Curtis dissimilarity matrices using a custom implementation based on Anderson’s distance-based partitioning (999 permutations, random seed = 42). Region and depth category were included as independent one-factor tests.

We also performed a complementary analysis without rarefaction for examining the impact of normalisation on the results. Count tables were Hellinger-transformed before calculating Bray–Curtis dissimilarities (Fig. S2).

Furthermore, the associations between sediment physical properties and eukaryotic community composition were examined. Each DNA sample was assigned the mean porosity within ±2.5 cm of its depth in the same core, approximately matching the 5-cm subsample thickness. Otherwise, porosity was linearly interpolated between adjacent measurements or, beyond the measured range, taken from the nearest measurement (24, 18 and 4 of 46 samples, respectively). Depth-adjusted porosity values (porosity residuals; see Section 2.6) were used in subsequent analyses. Read counts of zOTUs assigned to the same class were summed to generate class-level tables. Read counts were converted to relative abundance by dividing by the total number of reads per sample. These data formed the basis for the analyses described in Section 2.6.

### 2.6. Depth–porosity modelling and statistical analyses

To account for the general decrease in sediment porosity with burial depth, we first modelled the depth-porosity relationship for each coring site. High-resolution porosity profiles (Table S1; n = 1,195 measurements across eight holes) were fitted using two candidate models: a linear model and an exponential model.

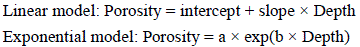

Model selection was based on the Akaike Information Criterion (AIC). For each hole, ΔAIC was calculated as AIC(linear) − AIC(exponential). Following Burnham and Anderson (2002), models differing by less than two AIC units were considered to have equivalent support, in which case the simpler linear model was retained. As a check, the community– and class-level analyses were repeated with exponential models and with the lowest-AIC model at each hole.

Porosity residuals (assigned porosity minus the model-predicted porosity at the sample depth) were used as depth-adjusted porosity. In the primary analysis, class-level relative abundances were converted to Bray–Curtis dissimilarities, and distance-based linear models (McArdle and Anderson, 2001) were fitted with site (eight sampling holes, as a factor), burial depth and log(burial depth + 1) as covariates (reduced model), and with depth-adjusted porosity added (full model). The log term accommodates near-surface non-linearity. Porosity effects (partial R², pseudo-F) were tested by permuting porosity residuals within sites (9,999 permutations). We pre-specified that porosity contributes at the community level if p < 0.05 for both markers. Sensitivity was tested by excluding the upper 5 cm and by omitting each site in turn. Variance partitioning based on adjusted R² (Peres-Neto et al., 2006) partitioned compositional variation among site, burial depth (linear and log terms) and porosity.

As a secondary, exploratory analysis, partial correlations between depth-adjusted porosity and the arcsine square-root transformed relative abundance of each class were calculated, controlling for burial depth and log(burial depth + 1). Classes with a mean relative abundance below 1 × 10□³ or occurring in fewer than five samples were excluded, and p-values were converted to Benjamini–Hochberg q-values (Benjamini and Hochberg, 1995). For classes with q < 0.05, post hoc robustness checks excluded the upper 5 cm, site U5, or each site in turn. Pearson correlations between relative abundance and burial depth were calculated for comparison. Statistical analyses and visualisations were performed in Python 3.13 using pandas 3.0.5, NumPy 2.5.3, SciPy 1.18.1 and matplotlib 3.11.2.

## 3. Results

### 3.1. Sequencing output and general composition

Sequencing of the 18S rRNA and COI genes from the Uranouchi Bay sediment samples yielded enough reads to detect taxonomic diversity at each site (18S rRNA: 14 million reads; COI: 6 million reads) (Fig. S1). For the 18S rRNA gene, most reads came from non-eukaryotic DNA, such as that of Bacteria and Archaea. The proportion of eukaryotic reads varied widely, from 0.1% to 68.5%, with a median of 3.6% (Table S2). Library construction failed for two 18S rRNA samples (U4, 10 cm; U5, 200 cm). Therefore, 44 and 46 samples were analysed for 18S rRNA and COI, respectively. After paired-end merging, quality filtering, chimera removal and denoising, 3,556 and 9,589 zOTUs were obtained for 18S and COI, respectively. All subsequent analyses used relative abundances based on the extracted eukaryotic reads.

### 3.2. Ordination overview of beta diversity

Principal coordinate analysis (PCoA) based on Bray–Curtis dissimilarity showed that, in both datasets, the cores from the inner bay (UUWs3261/4065/5361/6789) occupied a different region of the ordination space from the cores from the bay mouth (U3/U4/U5). This indicated that the sediment eukaryotic communities clearly differed between the inner bay and the bay mouth (Fig. 2). PERMANOVA further showed that, in both datasets, depth category as well as area significantly explained community composition (all p = 0.001; 999 permutations). The proportion of variation in community composition explained by area was larger in the 18S dataset (R² = 0.16, pseudo-F = 5.78) than in the COI dataset (R² = 0.07, pseudo-F = 3.18). Depth category also explained a significant proportion of the variation in community composition (18S: R² = 0.11, pseudo-F = 3.59; COI: R² = 0.08, pseudo-F = 3.83).

**Fig. 2.**
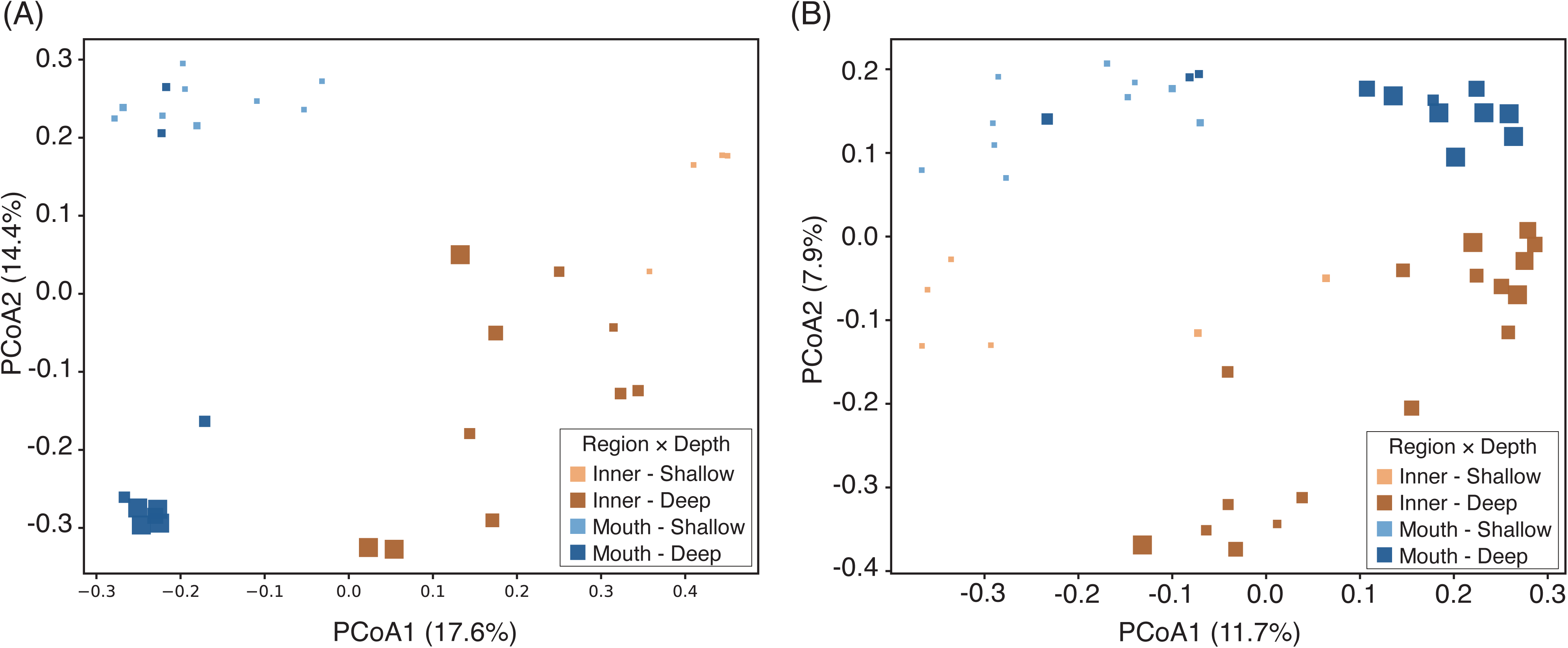
Principal coordinates analysis (PCoA) of eukaryotic community composition in the Uranouchi Bay sediments. (A) 18S rRNA gene; (B) COI gene. Ordinations are based on Bray–Curtis dissimilarities of Hellinger-transformed community composition derived from zOTU. Each sample was rarefied to 5,000 reads. Symbol size is proportional to sampling depth (cm). Colours indicate Region × Depth categories: Inner–Shallow (light orange), Inner–Deep (dark orange), Mouth–Shallow (light blue), and Mouth–Deep (dark blue), where Shallow = ≤50 cm and Deep = >50 cm sediment depth. The ordinations show a clear spatial separation between inner– and mouth-bay communities, with additional differentiation between shallow and deep horizons. Percent variance explained by the first two axes is indicated in parentheses.

Community composition also changed with depth. Within each area, deeper samples (larger symbols in Fig. 2) were separated from shallower samples. In the 18S dataset, this depth-related change was seen mainly along PCoA axis 2 in both areas. In contrast, in the COI dataset, the depth-related change was seen mainly along PCoA axis 1, and samples were separated by location within the bay along PCoA axis 2.

Ordination of Hellinger-transformed data without rarefaction reproduced both the separation between the inner bay and the bay mouth and the depth-related change in community composition (Fig. S2). This confirmed that the spatial structure we observed in community composition was not an artefact of differences in library size.

### 3.3. Spatial and vertical patterns of community composition

The eukaryotic community composition in Uranouchi Bay sediments differed between the inner bay and the bay mouth, with clear vertical differences within the cores (Figs. 2–4). The inner bay sites consistently showed strong terrestrial signals. The proportion of DNA from terrestrial plants (*Phragmoplastophyta*) was high, frequently exceeding 40% in deeper horizons (> 100 cm; Fig. 3). The inner-bay cores also showed a consistent presence of cyst-forming dinoflagellates (e.g., *Scrippsiella* spp., Fig. S3).

**Fig. 3.**
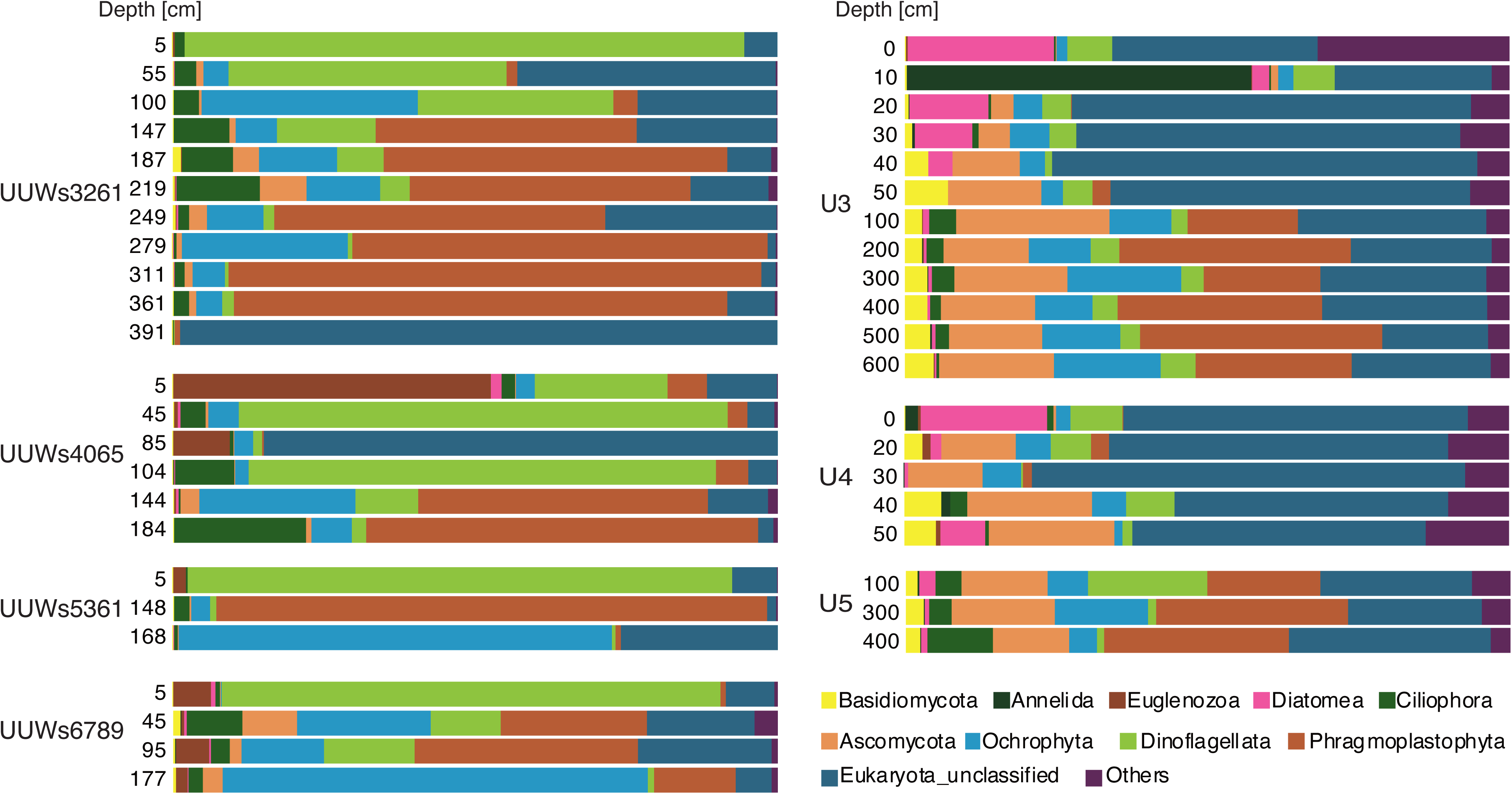
Phylum-level community composition of 18S rRNA gene sequences in the Uranouchi Bay sediments. Stacked bar charts show the relative abundances of major eukaryotic phyla detected across sediment depth intervals in each core. Left panels correspond to inner-bay sites (UUWs3261, UUWs4065, UUWs5361, UUWs6789), and right panels correspond to bay-mouth sites (U3, U4, U5, located near the entrance of the Uranouchi Bay). Depths are indicated in centimetres. Unassigned reads are shown as ‘Eukaryota_unclassified’, and minor groups are combined under ‘Others’. Relative abundances are based on raw counts after removal of non-eukaryotic sequences, without further rarefaction.

**Fig. 4.**
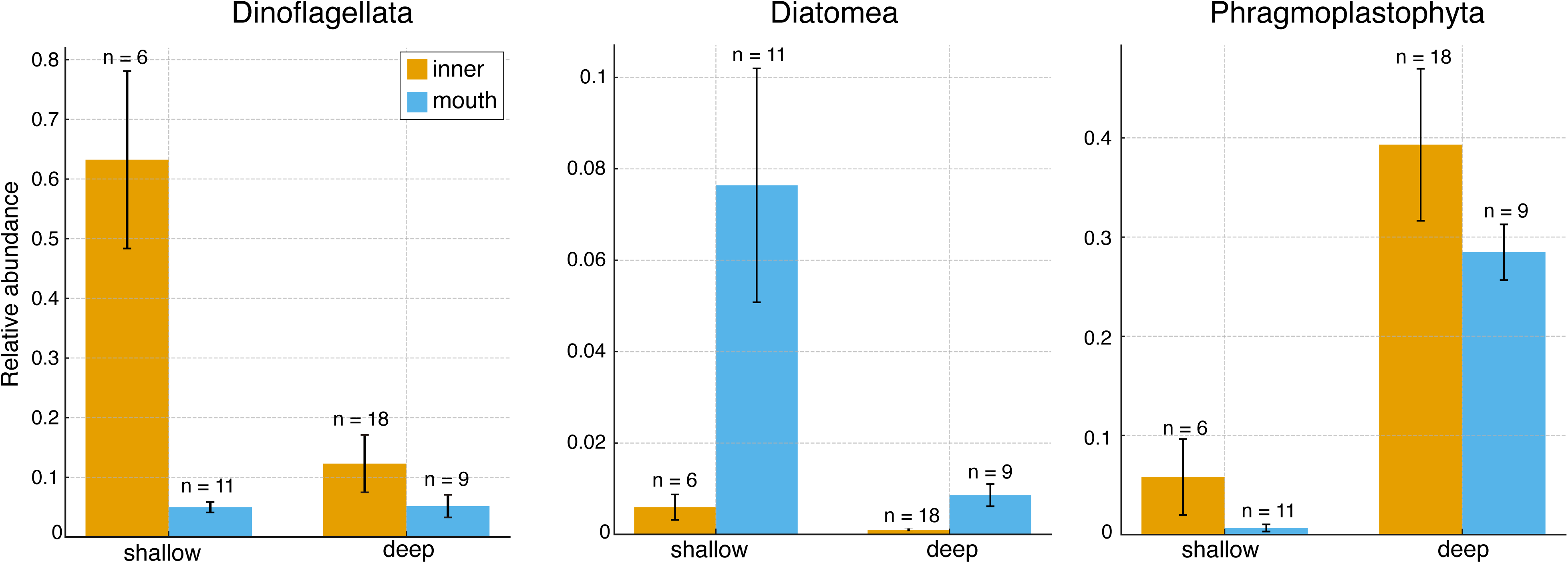
Relative abundances of key eukaryotic groups in the Uranouchi Bay sediments. Bar plots show the mean relative abundances (± standard error) of Dinoflagellata, Diatomea, and Phragmoplastophyta at shallow (0–50 cm) and deep (>50 cm) sediment layers in inner-bay and bay-mouth sites. Sample sizes (n) are indicated on each bar. Statistical significance was assessed using Mann–Whitney U tests; exact p-values are provided in Table S3.

In contrast, diatoms (Diatomea) were abundant at the bay-mouth sites (U3 and U4), particularly in the surface and upper layers (0–50 cm). To test these patterns, we compared the relative abundances of three representative taxa (Phragmoplastophyta, Diatomea and Dinoflagellata) between regions (inner bay vs. bay mouth) and between depth intervals (0–50 cm vs. > 50 cm) using Mann–Whitney U tests (Fig. 4; Supplementary Table S3). Dinoflagellata showed higher relative abundance in shallow than in deep inner-bay samples (p = 0.001) and in shallow inner-bay than in shallow mouth samples (p = 0.0002), consistent with their concentration in surface sediments of the restricted inner bay. In contrast, Diatomea were more abundant in deep mouth samples than in deep inner-bay samples (p = 0.0001), although they were also lower in deep than in shallow inner-bay samples (p = 0.030). Phragmoplastophyta were more abundant in shallow inner-bay than in shallow mouth samples (p = 0.049) and were higher in deep than in shallow samples in both regions (inner bay: p = 0.015; mouth: p = 0.0002).

### 3.4. Community composition of COI data

COI-based community composition also differed between the inner bay and the bay mouth (Fig. 5), although a large proportion of reads in most samples remained unassigned below the domain level (Eukaryota unclassified). Among assigned reads, Ascomycota were more prominent at the bay-mouth sites, particularly below 20 cm at U3 and in the deeper layers of U4 and U5, whereas their contribution was smaller in the inner-bay cores. Bacillariophyta were mainly detected in the surface layers of the bay-mouth sites (e.g., U3 at 0 cm, U4 at 0–10 cm, and U5 at 100 cm). Dinophyceae were detected mainly at the inner-bay sites, where they occurred in the uppermost samples (5 cm) of all four cores, whereas they were nearly absent from the bay-mouth sites. In addition to these general patterns, several individual samples were dominated by a single taxon. Bigyra accounted for a large fraction of reads at 100 and 391 cm in UUWs3261 and at 85 cm in UUWs4065, and Annelida dominated the 10 cm sample at U3. These spikes were largely confined to single horizons.

**Fig. 5.**
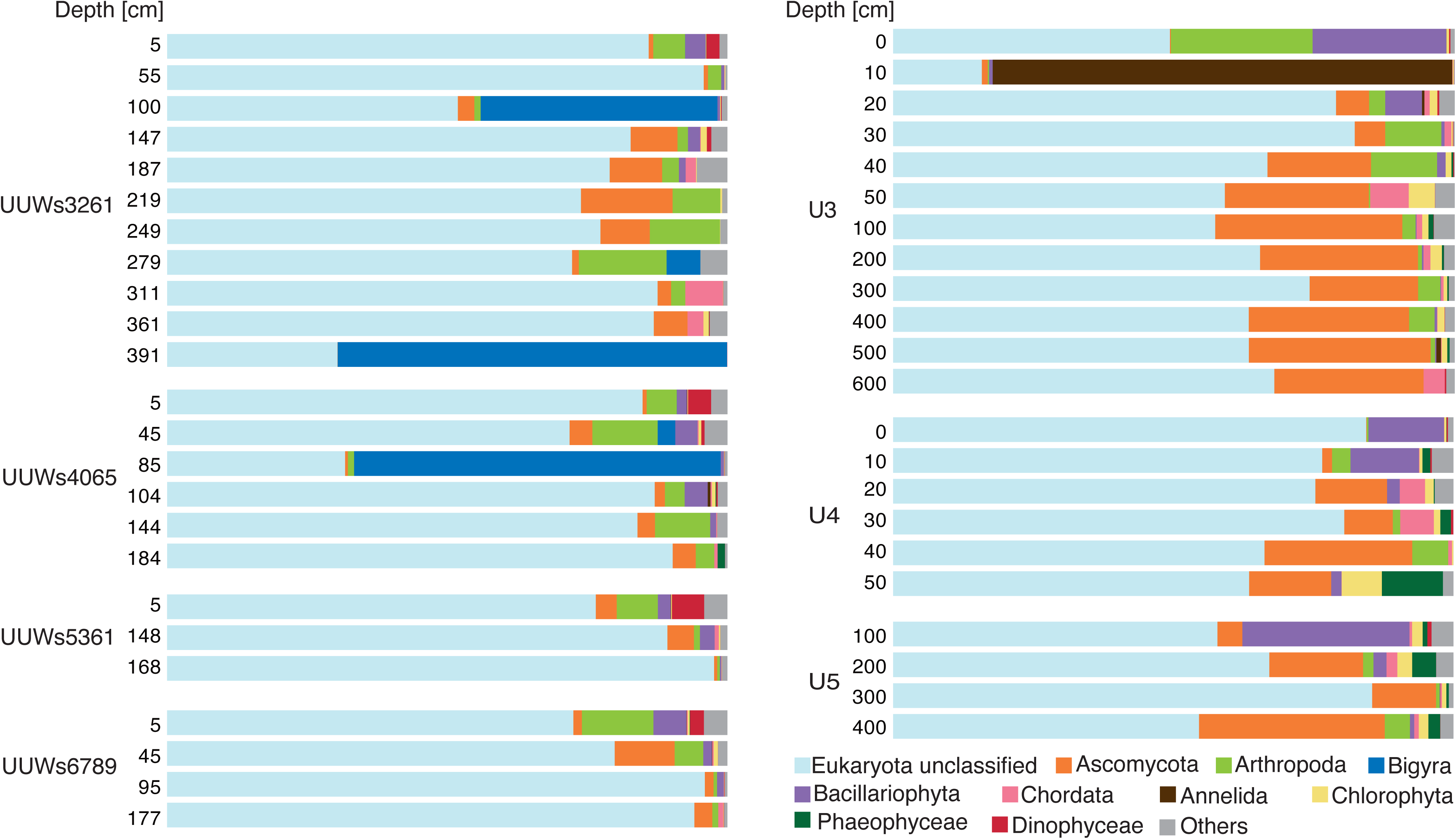
Community composition of COI gene sequences in the Uranouchi Bay sediments. Stacked bar charts show the relative abundances of major metazoan and protistan groups detected across sediment depth intervals in each core. Left panels correspond to inner-bay sites (UUWs3261, UUWs4065, UUWs5361, UUWs6789), and right panels correspond to bay-mouth sites (U3, U4, U5, located near the entrance of the Uranouchi Bay). Depths are indicated in centimetres. Unassigned reads are shown as ‘Eukaryota_unclassified’, and minor groups are combined under “Others”.

### 3.5. Relationship between sediment porosity and eDNA composition

Sediment porosity generally decreases linearly or exponentially with increasing depth (Bahr et al., 2001; Doan et al., 2023). We therefore fitted both models to the porosity depth profile of each of the eight cores. Site U3 was sampled twice, and each sampling was treated as an independent core. The linear model was clearly supported for U3_1 (ΔAIC = −3.9), whereas the two models showed little difference in fit for the remaining seven cores (|ΔAIC| < 2). The linear model was therefore adopted for all cores (Fig. S4; Table S4). Depth explained most of the variation in porosity in some cores (e.g. U3_1, R² = 0.95; UUWs6789, R² = 0.77) but had little explanatory power in others (U3_2, R² = 0.07; U5, R² < 0.01). In addition, in several cores the surface porosity was considerably higher than the value predicted by the linear model (Fig. S4).

Porosity corrected for depth using these models was then added as an explanatory variable. It did not significantly improve the explanation of eDNA composition for either 18S or COI (18S: partial R² = 1.0%, pseudo-F = 0.69, p = 0.74; COI: partial R² = 4.0%, pseudo-F = 2.11, p = 0.052). Surface porosity deviated from the models, so to remove its influence we repeated the analysis without samples from the upper 5 cm. This further reduced the contribution of porosity for COI (p = 0.37; 18S: p = 0.48). To exclude the influence of individual sites, the analysis was also repeated with one site omitted at a time. The resulting p values ranged from 0.38 to 0.92 for 18S and from 0.04 to 0.14 for COI (Table S5). In both the 18S and COI data sets, site explained the largest proportion of variation in eDNA composition (18S: 19.9%; COI: 8.2%), followed by sediment depth (4.9% and 2.7%, respectively). In contrast, the independent contribution of porosity was small (0.9% and 2.2%, respectively) (Fig. 6).

**Fig. 6.**
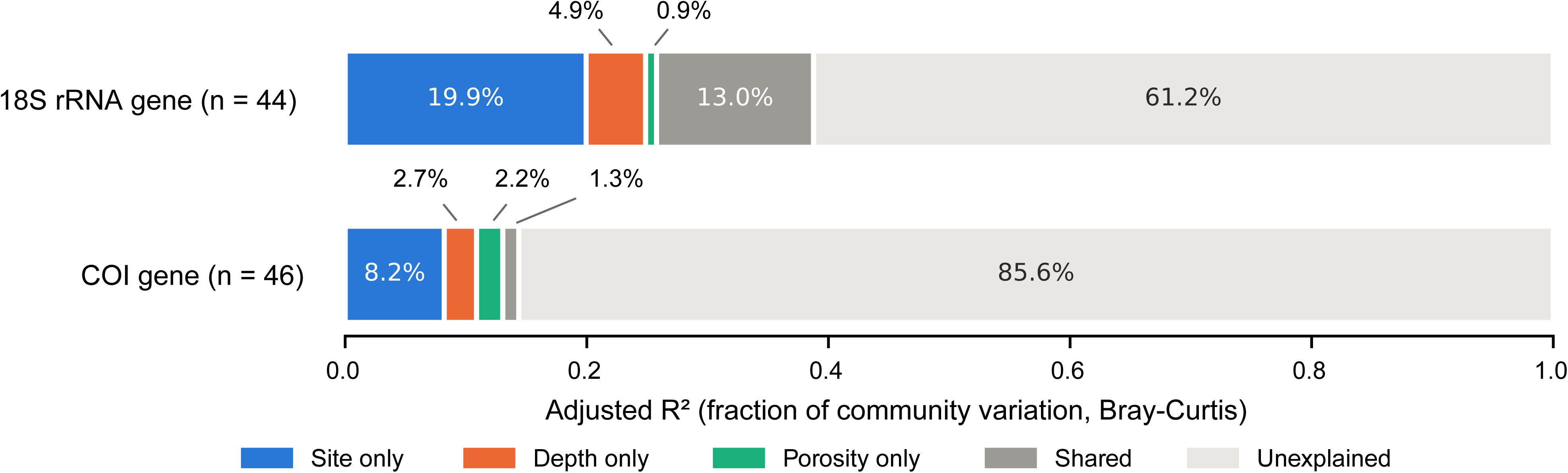
Variance partitioning of eukaryotic community composition (class-level Bray–Curtis dissimilarities) among site, burial depth (linear and log-transformed terms) and porosity for the 18S rRNA (n = 44) and COI (n = 46) datasets. Values are adjusted R² fractions; “Shared” combines all fractions shared among two or three factors. Porosity is depth-adjusted porosity (residuals from the depth–porosity model), calculated from interval-averaged values for each sample. The test for the contribution of depth-adjusted porosity beyond site and burial depth gave p = 0.74 (18S) and p = 0.052 (COI).

As a supplementary exploratory analysis, we calculated partial correlations at the class level. Depth-corrected porosity was significantly associated (q < 0.05) with 1 of 33 classes for 18S (Bacillariophyceae) and with 4 of 31 classes for COI (Coscinodiscophyceae, Bacillariophyceae, Dinophyceae and Branchiopoda) (Fig. S5). For both markers, diatom classes were negatively correlated with porosity (18S: Bacillariophyceae, r = −0.51, q = 0.012; COI: Coscinodiscophyceae, r = −0.62, q < 0.001; Bacillariophyceae, r = −0.56, q < 0.001). Mediophyceae in the 18S data set also showed a negative correlation, but it was not significant after correction for multiple comparisons (r = −0.42, p = 0.005, q = 0.07).

To determine whether these correlations depended on particular samples, we repeated the analysis twice: once without samples from the upper 5 cm, and once with one site omitted at a time. The negative correlations of diatom classes in COI were maintained in all cases (r = −0.42 and p ≤ 0.006 even in the weakest case). The correlation for 18S Bacillariophyceae was maintained when any single site was omitted. However, it was no longer observed when the upper 5 cm was excluded (r = −0.13, p = 0.44) (Fig. 7; Table S6). In contrast, the positive correlations of Dinophyceae and Branchiopoda in COI (both r ≈ +0.45, q = 0.015) depended on the surface samples. When the upper 5 cm was excluded, the direction of the correlation was reversed for Dinophyceae (r = −0.36, p = 0.025), and the correlation disappeared for Branchiopoda (r = +0.07, p = 0.67).

**Fig. 7.**
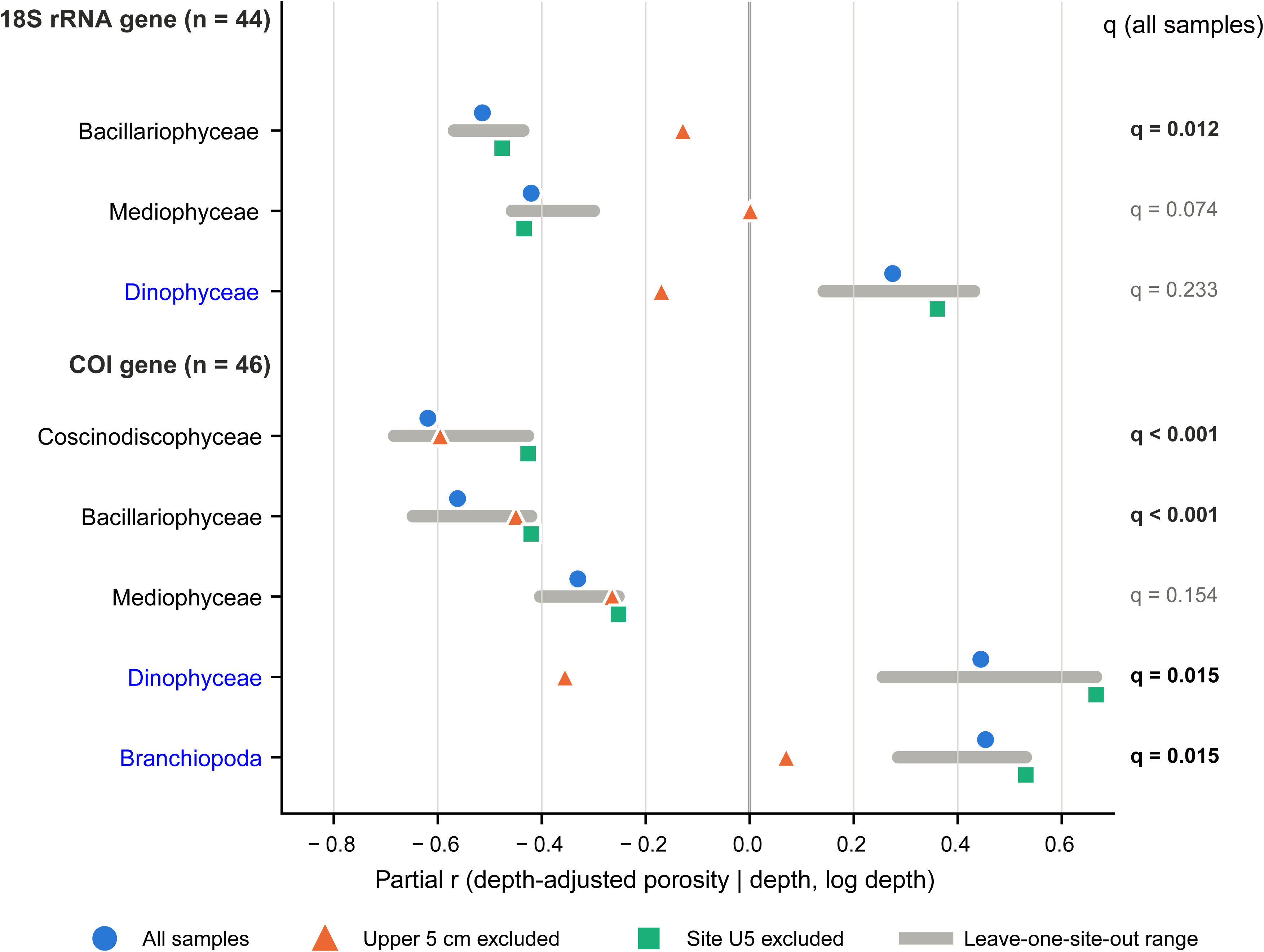
Robustness of class-level associations between depth-adjusted porosity and eukaryotic relative abundance (exploratory analysis). Partial correlation coefficients (controlling for burial depth and log-transformed burial depth) are shown for diatom classes and, for comparison, Dinophyceae and Branchiopoda, using all samples and after excluding samples from the upper 5 cm, excluding site U5, or omitting each site in turn (range). Robustness checks were performed post hoc.

We further examined whether the results were affected by how porosity was calculated. In the main analysis, porosity for each sample was calculated from the measurements within its sampling depth interval. When these values were replaced with the single measurement closest in depth, the correlations of diatom classes in COI became dependent on site U5 (Coscinodiscophyceae: r = −0.39 for all samples and r = −0.04 with U5 excluded). This was mainly because, at U5, the measurement closest to the 100 cm sample was located in a thin layer of low porosity (Fig. S6). With the interval-based approach used in the main analysis, the correlation was maintained when U5 was excluded (r = −0.43). In addition, the same five classes met q < 0.05 when the depth-correction model was replaced with the exponential model or with the model that had the lowest AIC for each core (Table S7b).

## 4. Discussion

### 4.1. Spatial and vertical structuring of sedimentary eDNA

Our results indicate that both basin geography and sediment depth influence the sedimentary eDNA composition in Uranouchi Bay. The limited water exchange of the bay promotes the retention of terrestrial material and locally produced plankton, which likely explains the higher proportions of dinoflagellate and land-plant DNA in inner-bay surface sediments. The marked differences in community composition between cores collected only a few kilometres apart indicate the importance of considering depositional setting when comparing sedimentary DNA records from different sites.

The dominance of dinoflagellates and diatoms in surface assemblages resembles 18S V9 records from another semi-enclosed Japanese bay (Segawa et al., 2022). The decline of diatoms and enrichment of land-plant DNA with depth are consistent with observations from lake sediments, where diatom DNA degrades much faster than vascular plant DNA, possibly because lignified cell walls physically shield plant biomolecules from post-burial degradation (Han et al., 2022). Another possibility is that the enrichment reflects past changes in terrestrial input, as plant DNA preservation in marine sediments has also been attributed primarily to pre-burial factors such as runoff and mineral supply (Herzschuh et al., 2025). Fungal DNA may similarly persist because of resistant cell walls or spores, although subsurface fungal activity cannot be excluded. Because obligate photosynthetic plankton and most zooplankton cannot survive after burial (Capo et al., 2021), eukaryotic DNA found in deeper layers is most plausibly interpreted as preserved DNA of organisms deposited in the past, with the possible exceptions of dormant stages and subsurface fungi (Torti et al., 2015). The persistence of *Scrippsiella*, a genus that forms calcareous resting cysts, below 100 cm in the inner bay, despite the overall decline of Dinoflagellata, illustrates the role of such dormant stages. In sediments of the Bay of Brest, most protist DNA was attributed to taxa capable of forming resting stages (Siano et al., 2021). *Scrippsiella* cysts retain their cell contents and germination capacity for several decades (Lundholm et al., 2011; Feifel et al., 2015), whereas relic DNA from vegetative cells degrades within months in sediments (Chai et al., 2024). Therefore, the *Scrippsiella* DNA detected in deeper layers more plausibly derives from buried cysts than from extracellular DNA, consistent with the role of semi-enclosed basins as seed banks for cyst-forming dinoflagellates.

COI provided additional information on metazoans and some protist lineages that are often underrepresented in sedimentary records (Gelabert et al., 2025). The Annelida-dominated surface sample at U3 most likely reflects living or recently dead polychaetes, whereas the Bigyra spikes in single deeper horizons may record the burial of DNA-rich detrital patches, as thraustochytrids and related lineages colonise organic aggregates (Wang et al., 2024). Taken together, these patterns are consistent with the widely recognised facts that sedimentary eDNA records do not provide unbiased information on past communities but are influenced by depositional setting and by traits that allow DNA to persist, so the absence of a taxon at depth should not be taken as evidence of its historical absence.

### 4.2. Sediment physical properties and implications for eDNA interpretation

In this study, we tested the hypothesis that sediment porosity explains variation in sedimentary eDNA composition beyond site and burial depth. At the community level, our results did not support this hypothesis. Porosity contributed only 1–4% of compositional variation when site and depth were accounted for, and this contribution was not significant under the pre-specified criterion. Furthermore, in the 18S dataset, the variation explained by porosity overlapped substantially with that explained by burial depth. Within the porosity range of these sediments (approximately 0.5–0.8), sedimentary eDNA composition therefore appears to be shaped mainly by depositional setting and burial depth. This result does not exclude effects of porosity outside this range or at spatial scales finer than our sampling resolution, but it indicates that porosity may not be a major confounder when sedimentary eDNA is compared among cores within a basin such as Uranouchi Bay.

However, we found a negative association between depth-adjusted porosity and diatom eDNA, which was observed in both markers and was robust in the COI dataset (Fig. 7; Fig. S7). Diatom DNA may be relatively better retained in more compacted layers, for example through association with siliceous frustules or adsorption to mineral surfaces, which can protect DNA from enzymatic degradation (Pietramellara et al., 2009; Sand et al., 2024), or diatom-rich material may be deposited preferentially in layers that differ in grain size or packing. Because this association emerged from exploratory analyses, it should be tested further together with other sediment properties such as grain size and organic carbon.

Our analyses also identify two practical pitfalls when linking sediment properties to sedimentary eDNA. One is that near-surface sediments combine high porosity with abundant DNA from recently deposited or living organisms (Torti et al., 2018), so taxon-level correlations can arise from the surface layer alone. In this study, for example, the positive Dinophyceae association disappeared or reversed when the upper 5 cm were excluded. The other is that assigning each eDNA sample a single nearest porosity value, rather than averaging over the sampled interval, can make results depend on thin layers or on arbitrary choices between equidistant measurements (Fig. S6). Porosity also covaries with grain size, mineralogy and diagenetic state, which can all affect how DNA associates with sediment particles (Sand et al., 2024). The porosity residuals used here therefore reflect several physical properties of the sediment together, and their association with eDNA does not necessarily indicate a direct effect of porosity.

### 4.3. Limitations and methodological considerations

In this study, burial depth was treated as a physical parameter rather than as a proxy for time, because no independent chronology was available. Dating by ²¹□Pb or ¹□ C would allow the depth– and porosity-related patterns to be interpreted on known timescales. Incomplete reference databases also constrained interpretation, leaving many sequences unassigned at lower taxonomic ranks and possibly obscuring finer-scale ecological signals. In addition, the attribution of the deep dinoflagellate signals to resting cysts and of the layer-specific Bigyra peaks to discrete depositional inputs remains only indicative, because intracellular and extracellular DNA were not separated.

The relationships between porosity and eDNA are correlative and cannot separate an effect of porosity from those of covarying sediment properties. The exploratory diatom association should be tested further in independent cores with measurements of grain size, biogenic silica, organic carbon and redox conditions, as well as through experiments on DNA retention in sediments of contrasting packing.

Eukaryote-targeting primers amplified substantial amounts of prokaryotic DNA, a common problem in sediment metabarcoding (Stoeck et al., 2010; Holman et al., 2025), and the eukaryotic fraction varied widely among samples (median 3.6%, range 0.1–68.5%; Table S2). All community analyses used relative abundances calculated within this eukaryotic fraction, so non-eukaryotic reads did not contribute directly to compositional estimates, although variable amplification may still have affected detection sensitivity. Although marker-specific biases cannot be excluded, the broad agreement between the 18S and COI datasets supports the main spatial and vertical patterns, as differences in community composition among samples are generally robust to methodological choices (Holman et al., 2025).

## 5. Conclusion

In this study, we examine the effect of porosity on eDNA preservation in marine sediment, and show that the eDNA composition is shaped mainly by depositional setting and burial depth, whereas sediment porosity plays a minor role. The eDNA communities from the inner bay and the bay mouth were different, and changed with depth in a taxon-dependent manner. Land-plant DNA and cyst-forming dinoflagellates persisted in deeper layers, while diatom DNA declined. Depth-adjusted porosity had little explanatory power for additional variation beyond site and burial depth. These results indicate that sedimentary eDNA in semi-enclosed bays is a selective archive of past communities, and that the influence of sediment physical properties should be tested rather than assumed. However, the association of porosity with diatom eDNA suggests that sediment physical properties may still affect the preservation of particular taxa. Future studies integrating porosity with grain size, geochemical indicators and other physical parameters will help understand the mechanisms by which sediment properties influence eDNA preservation.

## CRediT authorship contribution statement

**Tatsuhiko Hoshino:** Conceptualisation, Methodology, Investigation, Writing – review & editing, Writing – original draft, Funding acquisition. **Saki Toyama:** Investigation, Writing – review & editing. **Wataru Tanikawa:** Writing – review & editing, Investigation, Resources, Funding acquisition. **Masafumi Murayama**: Resources, Project administration, Funding acquisition, Supervision, Writing – review & editing.

## Declaration of competing interest

The authors declare that they have no known competing financial interests or personal relationships that could have appeared to influence the work reported in this paper.

## Declaration of generative AI and AI-assisted technologies in the manuscript preparation process

During the preparation of this work, the authors used ChatGPT and Claude (Anthropic) to assist with the generation and refinement of computational analysis scripts, as well as with the translation of portions of the manuscript from Japanese to English and language editing for clarity and readability. After using these tools, the authors carefully reviewed, validated, and edited all outputs as needed and take full responsibility for the content of the published article.

## Acknowledgements

The authors are grateful for the technical assistance provided by the staff working at Kochi Core Center. This work was partially supported by the Japan Society for the Promotion of Science (JSPS) Grant-in-Aid for Scientific Research JP21K19876 (to T.H.), JP23K22618 (to T.H.), JP20H04309 (to M.M., T.H., and W.T.), and JP16H03103 (to W.T. and M.M.)

## Data availability

The DNA sequence data reported in this study have been deposited in the DDBJ/ENA/GenBank databases under the accession number PRJDB37705.

## Notes

### Competing Interest Statement

The authors have declared no competing interest.

### Summary of Updates

We have revised the title to Is sediment porosity a confounder in sedimentary DNA records? A test in a temperate semi-enclosed bay. The Abstract, Highlights, Introduction, Discussion and Conclusions have been rewritten. We no longer treat porosity as an established factor in eDNA preservation; it is now presented as a hypothesis to be tested. We repeated the porosity analysis. Each DNA sample was assigned the mean porosity within 5 cm of its sampling depth, instead of the single nearest measurement. For each hole, linear and exponential depth-porosity models were compared using AIC, and the linear model was selected for all holes. The contribution of depth-corrected porosity was tested using distance-based linear models that included site, burial depth and log-transformed depth, with permutations restricted within sites. We also evaluated it by variance partitioning. After accounting for site and burial depth, porosity explained 1.0% (18S, p = 0.74) and 4.0% (COI, p = 0.052) of the variation in community composition, and neither contribution was significant. The phylum-level Spearman correlations and taxonomic groupings in version 1 have been replaced with an exploratory class-level partial correlation analysis with Benjamini-Hochberg correction. Diatom classes were negatively associated with depth-corrected porosity for both markers. The positive association of Dinophyceae reported in version 1 depended on near-surface samples, and we concluded that it is not robust. We added four sensitivity analyses: (1) excluding the upper 5 cm, (2) excluding one site at a time, (3) using an alternative depth-porosity model, and (4) assigning the nearest single porosity measurement. Figures 6 and 7 have been replaced (variance partitioning and the robustness of class-level associations). The supplementary figures and tables have been renumbered, and new ones have been added (porosity model fits, class-level partial correlations, the relationship between porosity and depth, the proportion of eukaryotic reads, and sensitivity analyses). The Methods now describe the uncertainty in porosity measurements, the sampling design at Site U3 (two holes), and our treatment of burial depth as a physical parameter rather than a proxy for age. We also added a new Limitations section and revised the text throughout for clarity.

## References

1. Armbrecht, L., Weber, M.E., Raymo, M.E., Peck, V.L., Williams, T., Warnock, J., Kato, Y., Hernández-Almeida, I., Hoem, F., Reilly, B., Hemming, S., Bailey, I., Martos, Y.M., Gutjahr, M., Percuoco, V., Allen, C., Brachfeld, S., Cardillo, F.G., Du, Z., Fauth, G., Fogwill, C., Garcia, M., Glüder, A., Guitard, M., Hwang, J.-H., Iizuka, M., Kenlee, B., O’Connell, S., Pérez, L.F., Ronge, T.A., Seki, O., Tauxe, L., Tripathi, S., Zheng, X., 2022. Ancient marine sediment DNA reveals diatom transition in Antarctica. Nat Commun 13, 5787. 10.1038/s41467-022-33494-4

2. Armbrecht, L.H., Coolen, M.J.L., Lejzerowicz, F., George, S.C., Negandhi, K., Suzuki, Y., Young, J., Foster, N.R., Armand, L.K., Cooper, A., Ostrowski, M., Focardi, A., Stat, M., Moreau, J.W., Weyrich, L.S., 2019. Ancient DNA from marine sediments: Precautions and considerations for seafloor coring, sample handling and data generation. Earth-Sci Rev 196, 102887. 10.1016/j.earscirev.2019.102887

3. Bahr, D.B., Hutton, E.W.H., Syvitski, J.P.M., Pratson, L.F., 2001. Exponential approximations to compacted sediment porosity profiles. Comput Geosci 27, 691–700. 10.1016/S0098-3004(00)00140-0

4. Benjamini, Y., Hochberg, Y., 1995. Controlling the false discovery rate: a practical and powerful approach to multiple testing. J. R. Stat. Soc. Ser. B 57, 289–300.

5. Blum, P., 1997. Physical properties handbook: a guide to the shipboard measurement of physical properties of deep-sea cores. Ocean Drill. Progr. Tech. Note 26. 10.2973/odp.tn.26.1997.

6. Boere, A.C., Rijpstra, W.I.C., De Lange, G.J., Sinninghe Damsté, J.S., Coolen, M.J.L., 2011. Preservation potential of ancient plankton DNA in Pleistocene marine sediments. Geobiology 9, 377–393. 10.1111/j.1472-4669.2011.00290.x

7. Burnham, K.P., Anderson, D.R., 2002. Model Selection and Multimodel Inference: A Practical Information-Theoretic Approach, 2nd ed. Springer, New York. 10.1007/b97636

8. Campbell, M.A., Ward, I., Blyth, A., Allentoft, M.E., 2025. Using sedimentary ancient DNA in coastal and marine contexts to explore past human–environmental interactions in Australia. Phil. Trans. R. Soc. B 380, 20240032. 10.1098/rstb.2024.0032

9. Capo, E., Giguet-Covex, C., Rouillard, A., Nota, K., Heintzman, P.D., Vuillemin, A., Ariztegui, D., Arnaud, F., Belle, S., Bertilsson, S., Bigler, C., Bindler, R., Brown, A.G., Clarke, C.L., Crump, S.E., Debroas, D., Englund, G., Ficetola, G.F., Garner, R.E., Gauthier, J., Gregory-Eaves, I., Heinecke, L., Herzschuh, U., Ibrahim, A., Kisand, V., Kjær, K.H., Lammers, Y., Littlefair, J., Messager, E., Monchamp, M.-E., Olajos, F., Orsi, W., Pedersen, M.W., Rijal, D.P., Rydberg, J., Spanbauer, T., Stoof-Leichsenring, K.R., Taberlet, P., Talas, L., Thomas, C., Walsh, D.A., Wang, Y., Willerslev, E., Van Woerkom, A., Zimmermann, H.H., Coolen, M.J.L., Epp, L.S., Domaizon, I., G. Alsos, I., Parducci, L., 2021. Lake Sedimentary DNA Research on Past Terrestrial and Aquatic Biodiversity: Overview and Recommendations. Quaternary 4, 6. 10.3390/quat4010006

10. Chai, Z., Liu, Y., Jia, S., Li, F., Hu, Z., Deng, Y., Yue, C., Tang, Y.Z., 2024. DNA and RNA stability of marine microalgae in cold-stored sediments and its implications in metabarcoding analyses. Int J Mol Sci 25, 1724. 10.3390/ijms25031724

11. Corinaldesi, C., Barucca, M., Luna, G.M., Dell’Anno, A., 2011. Preservation, origin and genetic imprint of extracellular DNA in permanently anoxic deep-sea sediments. Mol Ecol 20, 642–654. 10.1111/j.1365-294X.2010.04958.x

12. Crecchio, C., Stotzky, G., 1998. Binding of DNA on humic acids: Effect on transformation of Bacillus subtilis and resistance to DNase. Soil Biol Biochem 30, 1061–1067. 10.1016/S0038-0717(97)00248-4

13. De Schepper, S., Ray, J.L., Skaar, K.S., Sadatzki, H., Ijaz, U.Z., Stein, R., Larsen, A., 2019. The potential of sedimentary ancient DNA for reconstructing past sea ice evolution. ISME J 13, 2566–2577. 10.1038/s41396-019-0457-1

14. Deiner, K., Bik, H.M., Mächler, E., Seymour, M., Lacoursière Roussel, A., Altermatt, F., Creer, S., Bista, I., Lodge, D.M., De Vere, N., Pfrender, M.E., Bernatchez, L., 2017. Environmental DNA metabarcoding: Transforming how we survey animal and plant communities. Mol Ecol 26, 5872–5895. 10.1111/mec.14350

15. Doan, M., Dutilleul, J., Henry, P., 2023. Effective porosity profile at IODP site C0002 in the heart of the Nankai Accretionary Prism, and its use for predicting in situ seismic velocities. Geophysic Res Lett 50, e2022GL100209. 10.1029/2022GL100209

16. Edgar, R.C., 2016. UNOISE2: improved error-correction for Illumina 16S and ITS amplicon sequencing. bioRxiv 081257. 10.1101/081257

17. Feifel, K.M., Fletcher, S.J., Watson, L.R., Moore, S.K., Lessard, E.J., 2015. *Alexandrium* and *Scrippsiella* cyst viability and cytoplasmic fullness in a 60-cm sediment core from Sequim Bay, WA. Harmful Algae 47, 56–65. 10.1016/j.hal.2015.05.009

18. Funaki, H., Gaonkar, C.C., Nishimura, T., Tanaka, K., Kamimura, K., Kaji, T., Nagasaki, K., Adachi, M., 2025. Vertical distribution of harmful algae in the sediment of Uranouchi Inlet by metabarcoding. Front. Protistol. 3, 1612811. 10.3389/frpro.2025.1612811

19. Gelabert, P., Oberreiter, V., Straus, L.G., Morales, M.R.G., Sawyer, S., Marín-Arroyo, A.B., Geiling, J.M., Exler, F., Brueck, F., Franz, S., Cano, F.T., Szedlacsek, S., Zelger, E., Hämmerle, M., Zagorc, B., Llanos-Lizcano, A., Cheronet, O., Tejero, J.-M., Rattei, T., Kraemer, S.M., Pinhasi, R., 2025. A sedimentary ancient DNA perspective on human and carnivore persistence through the Late Pleistocene in El Mirón Cave, Spain. Nat Commun 16, 107. 10.1038/s41467-024-55740-7

20. Gupta, L.P., Tanikawa, W., Hamada, Y., Hirose, T., Ahagon, N., Sugihara, T., Abe, N., Nomura, S., Masaki, Y., Wu, H.Y., Lin, W., Kinoshita, M., Yamada, Y., 2019. Examination of gas hydrate-bearing deep ocean sediments by X-ray Computed Tomography and verification of physical property measurements of sediments. Marine and Petroleum Geology 108, 239–248. 10.1016/j.marpetgeo.2018.05.033.

21. Han, X., Tolu, J., Deng, L., Fiskal, A., Schubert, C.J., Winkel, L.H.E., Lever, M.A., 2022. Long-term preservation of biomolecules in lake sediments: potential importance of physical shielding by recalcitrant cell walls. PNAS Nexus 1, pgac076. 10.1093/pnasnexus/pgac076

22. Herzschuh, U., Weiß, J.F., Stoof-Leichsenring, K.R., Harms, L., Nürnberg, D., Müller, J., 2025. Dynamic land-plant carbon sources in marine sediments inferred from ancient DNA. Commun Earth Environ 6, 78. 10.1038/s43247-025-02014-9

23. Holman, L.E., Zampirolo, G., Gyllencreutz, R., Scourse, J., Frøslev, T., Carøe, C., Gopalakrishnan, S., Pedersen, M.W., Bohmann, K., 2025. Navigating Past Oceans: Comparing Metabarcoding and Metagenomics of Marine Ancient Sediment Environmental DNA. Molecular Ecology Resources 25, e14086. 10.1111/1755-0998.14086

24. Hoshino, T., Inagaki, F., 2019. Abundance and distribution of Archaea in the subseafloor sedimentary biosphere. ISME J 13, 227–231. 10.1038/s41396-018-0253-3

25. Hoshino, T., Inagaki, F., 2024. Distribution of eukaryotic environmental DNA in global subseafloor sediments. Prog Earth Planet Sci 11, 19. 10.1186/s40645-024-00621-2

26. Huettel, M., Gust, G., 1992. Impact of bioroughness on interfacia solute exchange in permeable sediments. Mar. Ecol. Prog. Ser. 89, 253–267. 10.3354/meps089253

27. Lejzerowicz, F., Esling, P., Majewski, W., Szczuciński, W., Decelle, J., Obadia, C., Arbizu, P.M., Pawlowski, J., 2013. Ancient DNA complements microfossil record in deep-sea subsurface sediments. Biol. Lett. 9, 20130283. 10.1098/rsbl.2013.0283

28. Leray, M., Knowlton, N., Machida, R.J., 2022. MIDORI2: A collection of quality controlled, preformatted, and regularly updated reference databases for taxonomic assignment of eukaryotic mitochondrial sequences. Environ DNA 4, 894–907. 10.1002/edn3.303

29. Leray, M., Yang, J.Y., Meyer, C.P., Mills, S.C., Agudelo, N., Ranwez, V., Boehm, J.T., Machida, R.J., 2013. A new versatile primer set targeting a short fragment of the mitochondrial COI region for metabarcoding metazoan diversity: application for characterizing coral reef fish gut contents. Front Zool 10, 34. 10.1186/1742-9994-10-34

30. Lundholm, N., Ribeiro, S., Andersen, T.J., Koch, T., Godhe, A., Ekelund, F., Ellegaard, M., 2011. Buried alive – germination of up to a century-old marine protist resting stages. Phycologia 50, 629–640. 10.2216/11-16.1

31. McArdle, B.H., Anderson, M.J., 2001. Fitting multivariate models to community data: a comment on distance-based redundancy analysis. Ecology 82, 290–297.

32. Meyer, C.P., Geller, J.B., Paulay, G., 2005. Fine scale endemism on coral reefs: archipelagic differentiation in turbinid gastropods. Evol 59, 113. 10.1554/04-194

33. Nguyen, N., Pawłowska, J., Zajaczkowski, M., Weiner, A.K.M., Cordier, T., Grant, D.M., De Schepper, S., Pawłowski, J., 2024. Taxonomic and abundance biases affect the record of marine eukaryotic plankton communities in sediment DNA archives. Mol Ecol Resour 24, e14014. 10.1111/1755-0998.14014

34. Pawłowska, J., Lejzerowicz, F., Esling, P., Szczuciński, W., Zajączkowski, M., Pawlowski, J., 2014. Ancient DNA sheds new light on the Svalbard foraminiferal fossil record of the last millennium. Geobiology 12, 277–288. 10.1111/gbi.12087

35. Pawlowski, J., Kelly-Quinn, M., Altermatt, F., Apothéloz-Perret-Gentil, L., Beja, P., Boggero, A., Borja, A., Bouchez, A., Cordier, T., Domaizon, I., Feio, M.J., Filipe, A.F., Fornaroli, R., Graf, W., Herder, J., Van Der Hoorn, B., Iwan Jones, J., Sagova-Mareckova, M., Moritz, C., Barquín, J., Piggott, J.J., Pinna, M., Rimet, F., Rinkevich, B., Sousa-Santos, C., Specchia, V., Trobajo, R., Vasselon, V., Vitecek, S., Zimmerman, J., Weigand, A., Leese, F., Kahlert, M., 2018. The future of biotic indices in the ecogenomic era: Integrating (e)DNA metabarcoding in biological assessment of aquatic ecosystems. Sci Total Environ 637–638, 1295–1310. 10.1016/j.scitotenv.2018.05.002

36. Peres-Neto, P.R., Legendre, P., Dray, S., Borcard, D., 2006. Variation partitioning of species data matrices: estimation and comparison of fractions. Ecology 87, 2614–2625.

37. Pietramellara, G., Ascher, J., Borgogni, F., Ceccherini, M.T., Guerri, G., Nannipieri, P., 2009. Extracellular DNA in soil and sediment: fate and ecological relevance. Biol Fertil Soils 45, 219–235. 10.1007/s00374-008-0345-8

38. Quast, C., Pruesse, E., Yilmaz, P., Gerken, J., Schweer, T., Yarza, P., Peplies, J., Glöckner, F.O., 2012. The SILVA ribosomal RNA gene database project: improved data processing and web-based tools. Nucleic Acid Res 41, D590–D596. 10.1093/nar/gks1219

39. Sand, K.K., Jelavić, S., Kjær, K.H., Prohaska, A., 2024. Importance of EDNA taphonomy and sediment provenance for robust ecological inference: Insights from interfacial geochemistry. Environ DNA 6, e519. 10.1002/edn3.519

40. Schloss, P.D., Westcott, S.L., Ryabin, T., Hall, J.R., Hartmann, M., Hollister, E.B., Lesniewski, R.A., Oakley, B.B., Parks, D.H., Robinson, C.J., Sahl, J.W., Stres, B., Thallinger, G.G., Van Horn, D.J., Weber, C.F., 2009. Introducing mothur: Open-source, platform-independent, community-supported software for describing and comparing microbial communities. Appl Environ Microbiol 75, 7537–7541. 10.1128/AEM.01541-09

41. Segawa, Y., Yamamoto, M., Kuwae, M., Moriya, K., Suzuki, H., Suzuki, K., 2022. Reconstruction of the eukaryotic communities in Beppu Bay over the past 50 years based on sedimentary DNA barcoding. JGR Biogeosci 127, e2022JG006825. 10.1029/2022JG006825

42. Shiraki, K., Ito, M., Miyake, K., Yukihiro, M., 1996. Water quality survey of Uranouchi Bay (focusing on nitrogen and phosphorus). Annu. Rep. Kochi Prefect. Environ. Res. Cent. 13, 51–57 (in Japanese).

43. Siano, R., Lassudrie, M., Cuzin, P., Briant, N., Loizeau, V., Schmidt, S., Ehrhold, A., Mertens, K.N., Lambert, C., Quintric, L., Noël, C., Latimier, M., Quéré, J., Durand, P., Penaud, A., 2021. Sediment archives reveal irreversible shifts in plankton communities after World War II and agricultural pollution. Curr Biol 31, 2682–2689.e7. 10.1016/j.cub.2021.03.079

44. Stoeck, T., Bass, D., Nebel, M., Christen, R., Jones, M.D.M., Breiner, H.-W., Richards, T.A., 2010. Multiple marker parallel tag environmental DNA sequencing reveals a highly complex eukaryotic community in marine anoxic water. Mol Ecol 19, 21–31. 10.1111/j.1365-294X.2009.04480.x

45. Taberlet, P., Coissac, E., Hajibabaei, M., Rieseberg, L.H., 2012. Environmental DNA. Mol Ecol 21, 1789–1793. 10.1111/j.1365-294X.2012.05542.x

46. Torti, A., Lever, M.A., Jørgensen, B.B., 2015. Origin, dynamics, and implications of extracellular DNA pools in marine sediments. Marine Genomics 24, 185–196. 10.1016/j.margen.2015.08.007

47. Torti, A., Jørgensen, B.B., Lever, M.A., 2018. Preservation of microbial DNA in marine sediments: insights from extracellular DNA pools. Environ Microbiol 20, 4526–4542. 10.1111/1462-2920.14401

48. Wang, Q., Zhang, Y., Hui, R., Zhu, Y., 2024. Marine thraustochytrid: exploration from taxonomic challenges to biotechnological applications. Front. Mar. Sci. 11, 1371713. 10.3389/fmars.2024.1371713

