## Supplemental Figure S1 for "Is sediment porosity a confounder in sedimentary DNA records? A test in a temperate semi-enclosed bay"

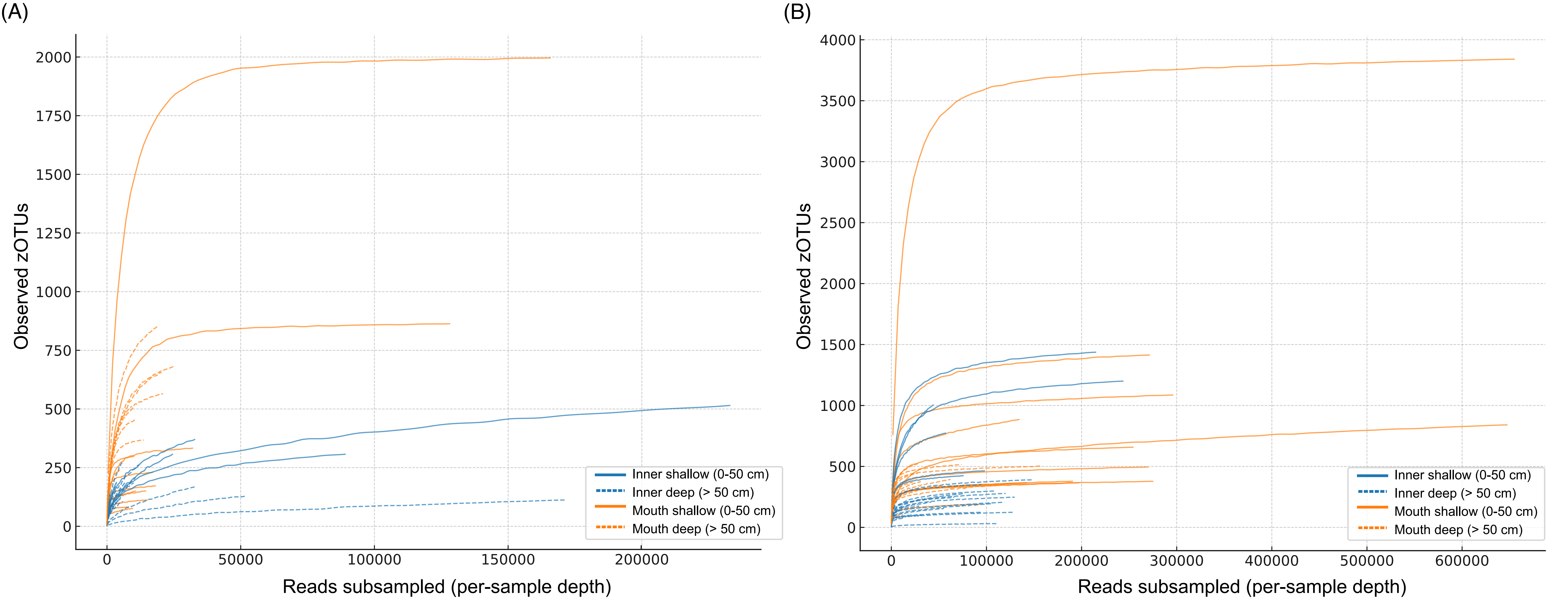


**FIGURE S1**

Rarefaction curves of (A) 18S rRNA gene (Eukaryota only) and (B) COI gene datasets, showing the accumulation of observed zOTUs as a function of sequencing depth for all sediment samples after removal of non-eukaryotic sequences. Each curve represents one sample and is drawn up to its maximum sequencing depth. Colors indicate sampling location (blue: inner bay; orange: mouth of the bay), and line styles indicate sediment depth (solid line: shallow, 0–50 cm; dashed line: deep, >50 cm). Curves for both markers reach saturation for most samples, demonstrating that sequencing depth was sufficient to capture the majority of eukaryotic diversity across samples. This justified inclusion of all samples in downstream analyses of relative abundances of major taxonomic groups such as terrestrial plants, diatoms, dinoflagellates, and metazoans.
