## Supplemental Figure S2 for "Is sediment porosity a confounder in sedimentary DNA records? A test in a temperate semi-enclosed bay"

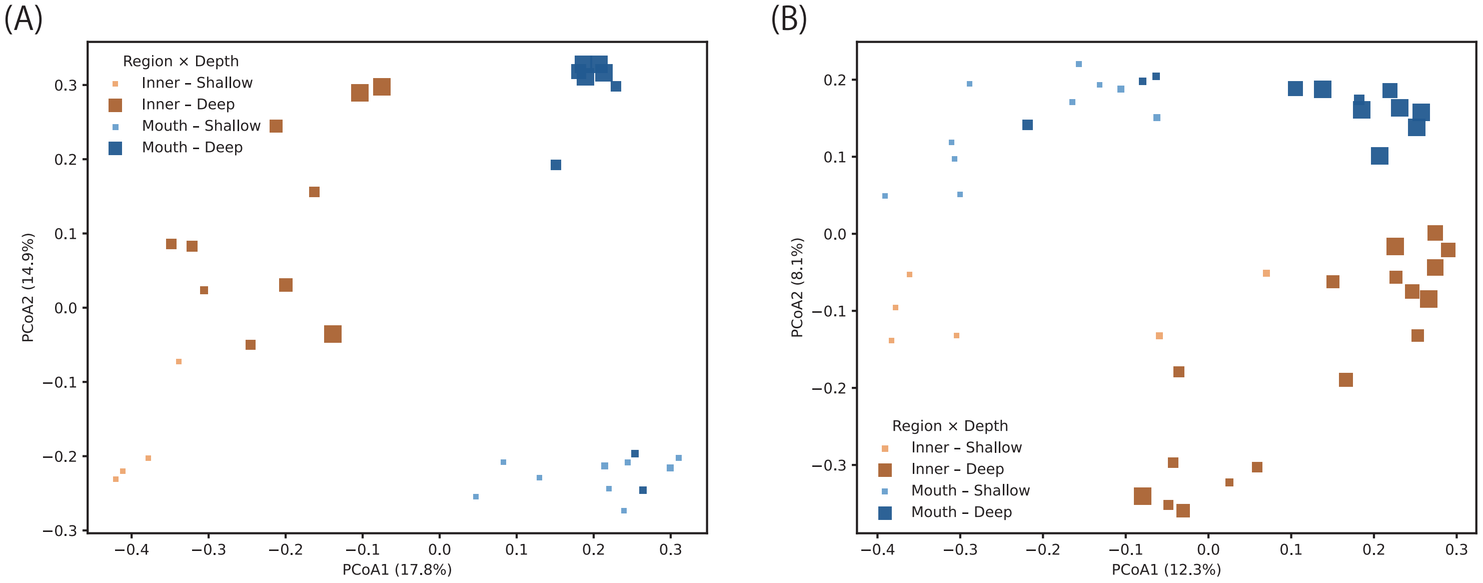


**FIGURE S2**

Principal coordinates analysis (PCoA) of eukaryotic community composition in Uranouchi Bay sediments. (A) 18S rRNA gene; (B) COI gene. Ordinations are based on Bray–Curtis dissimilarities of Hellinger-transformed community composition derived from zOTUs, using total sum scaling (TSS) without rarefaction. Symbol size is proportional to sampling depth (cm). Colors indicate Region × Depth categories: Inner–Shallow (light orange), Inner–Deep (dark orange), Mouth–Shallow (light blue), and Mouth–Deep (dark blue), where Shallow = <50 cm and Deep = >50 cm sediment depth. The ordinations confirm the clear separation between inner- and mouth-bay communities observed in the rarefied datasets (Figure 2), and demonstrate that the spatial patterns are robust to the choice of normalization method.
