## Supplemental Figure S3 for "Is sediment porosity a confounder in sedimentary DNA records? A test in a temperate semi-enclosed bay"

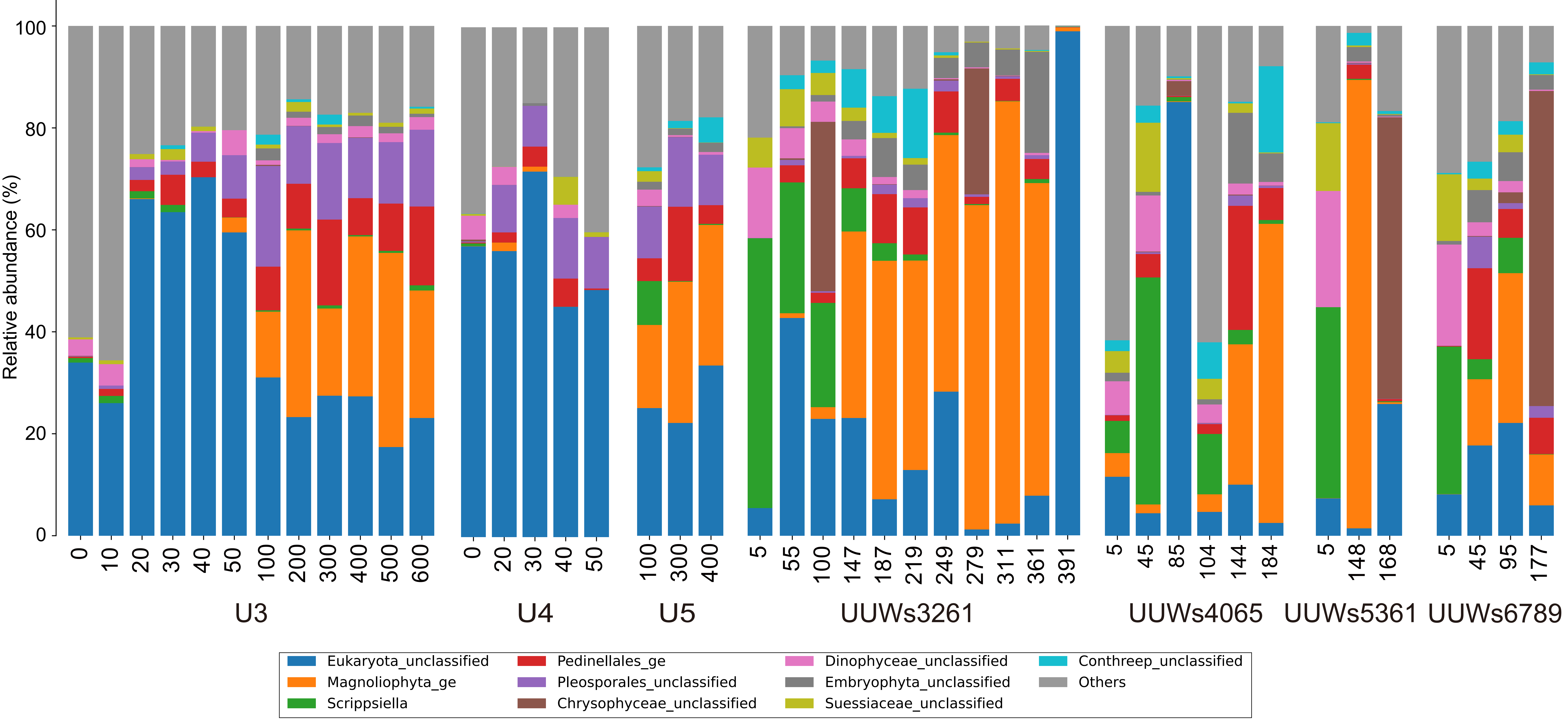


Fig. S3. Composition of the ten most abundant taxa (lowest assigned rank) in the 18S rRNA gene dataset, shown for each sample. Bars are grouped by site (inner bay: UUWs3261, UUWs4065, UUWs5361, UUWs6789; bay mouth: U3, U4, U5) and ordered by depth; sample labels give site and depth (cm). Relative abundances were calculated within the eukaryotic fraction. Taxa not assigned to genus level are labelled with the lowest assigned rank (suffix "_ge" or "_unclassified"); all remaining taxa are grouped as "Others". Note the persistence of Scrippsiella below 100 cm in the inner-bay cores.
