## Supplemental Figure S4 for "Is sediment porosity a confounder in sedimentary DNA records? A test in a temperate semi-enclosed bay"

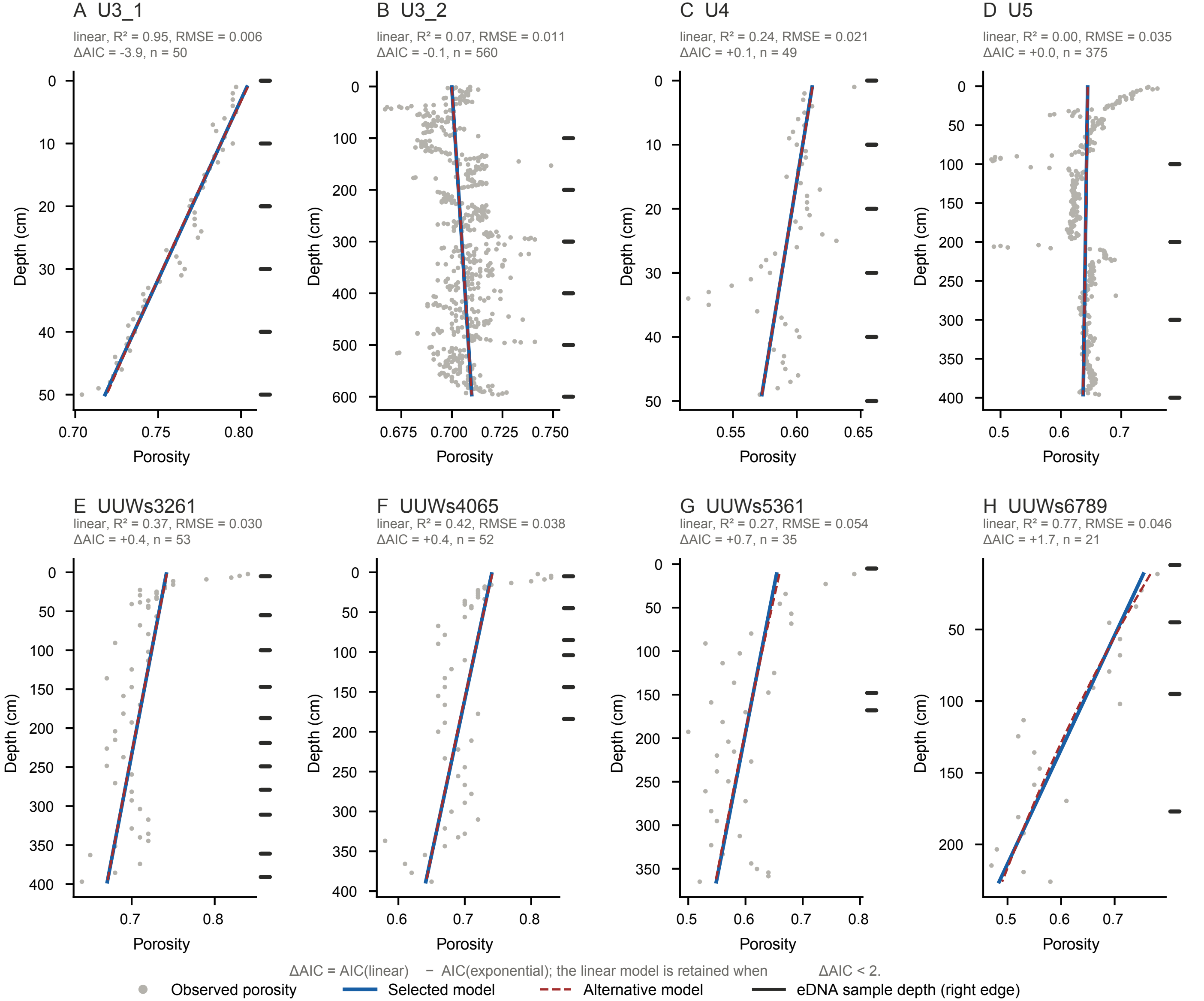


**Fig. S4.** Depth–porosity profiles and model fits for each sampling hole: (A) U3_1, (B) U3_2, (C) U4, (D) U5, (E) UUWs3261, (F) UUWs4065, (G) UUWs5361 and (H) UUWs6789. Grey dots show observed porosity, determined from X-ray CT data for the bay-mouth cores (A–D) and by gravimetric and pycnometric measurements for the inner-bay cores (E–H). Linear and exponential models were fitted to each profile. Solid blue lines indicate the selected model, and dashed red lines indicate the alternative model. ΔAIC was calculated as AIC(linear) − AIC(exponential), and the linear model was retained when ΔAIC < 2. The $R^2$, root mean square error (RMSE), ΔAIC and number of porosity measurements (n) of the selected model are given above each panel. Black marks on the right edge of each panel indicate the depths of the eDNA samples. Note that the depth and porosity axes differ among panels.
