## Supplemental Figure S5 for "Is sediment porosity a confounder in sedimentary DNA records? A test in a temperate semi-enclosed bay"

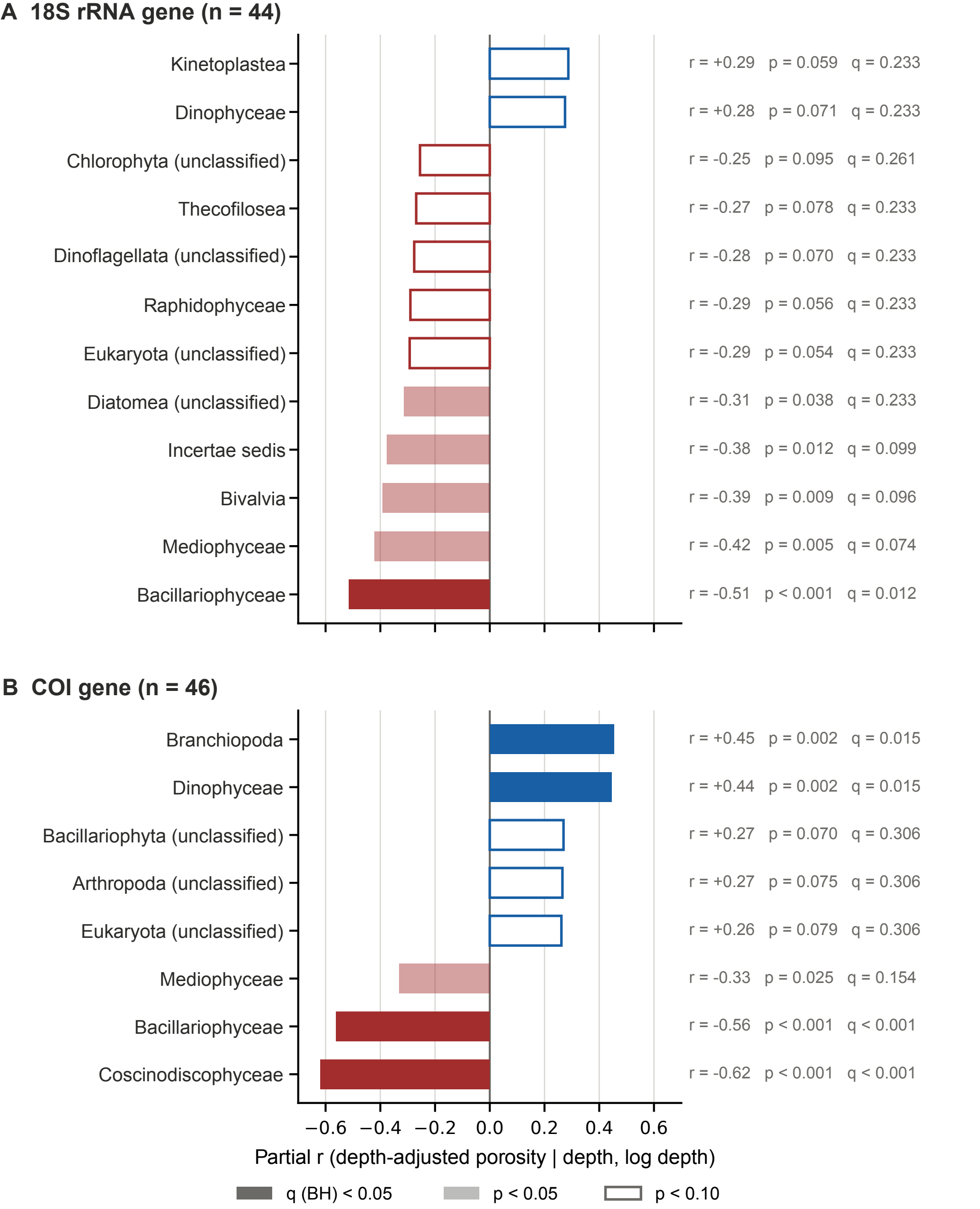


**Fig. S5.** Partial correlations between depth-adjusted porosity and the relative abundance of eukaryotic classes for (A) the 18S rRNA gene (n = 44) and (B) the COI gene (n = 46) datasets. Partial correlation coefficients were calculated between porosity residuals and arcsine square-root transformed relative abundance, controlling for burial depth and log(burial depth + 1). Classes with a mean relative abundance below $1 \times 10^{-3}$ or occurring in fewer than five samples were excluded, and only classes with $p < 0.10$ are shown. Red and blue bars indicate negative and positive correlations, respectively. Filled dark bars indicate Benjamini–Hochberg $q < 0.05$, filled light bars $p < 0.05$, and open bars $p < 0.10$. The correlation coefficient, p-value and q-value for each class are given on the right.
