## Supplemental Figure S6 for "Is sediment porosity a confounder in sedimentary DNA records? A test in a temperate semi-enclosed bay"

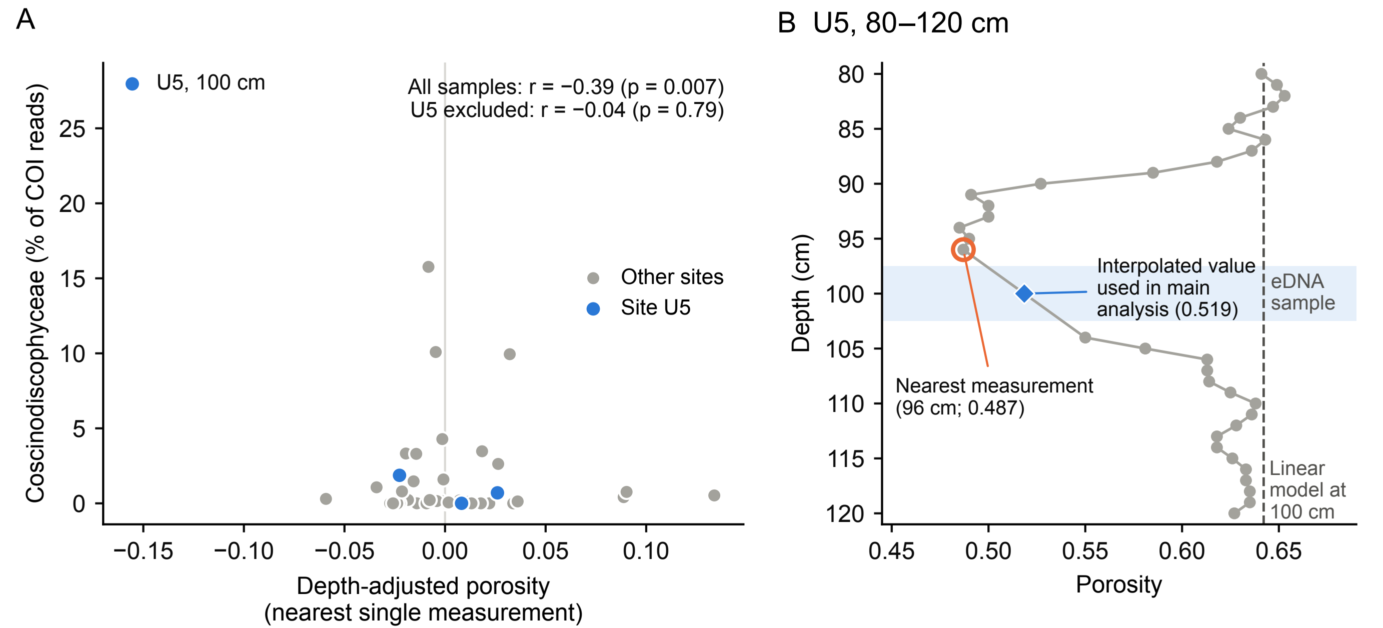


**Fig. S6.** Sensitivity of the diatom–porosity association to the porosity assignment method. (A) Relative abundance of Coscinodiscophyceae (COI) versus depth-adjusted porosity calculated from the single nearest porosity measurement to each sample. Partial correlations (controlling for burial depth) are shown for all samples and after excluding site U5. (B) Porosity profile of site U5 between 80 and 120 cm. The shaded band indicates the eDNA sample interval (97.5–102.5 cm); the open circle marks the nearest measurement (96 cm), which falls within a thin low-porosity
