## Supplemental Figure S7 for "Is sediment porosity a confounder in sedimentary DNA records? A test in a temperate semi-enclosed bay"

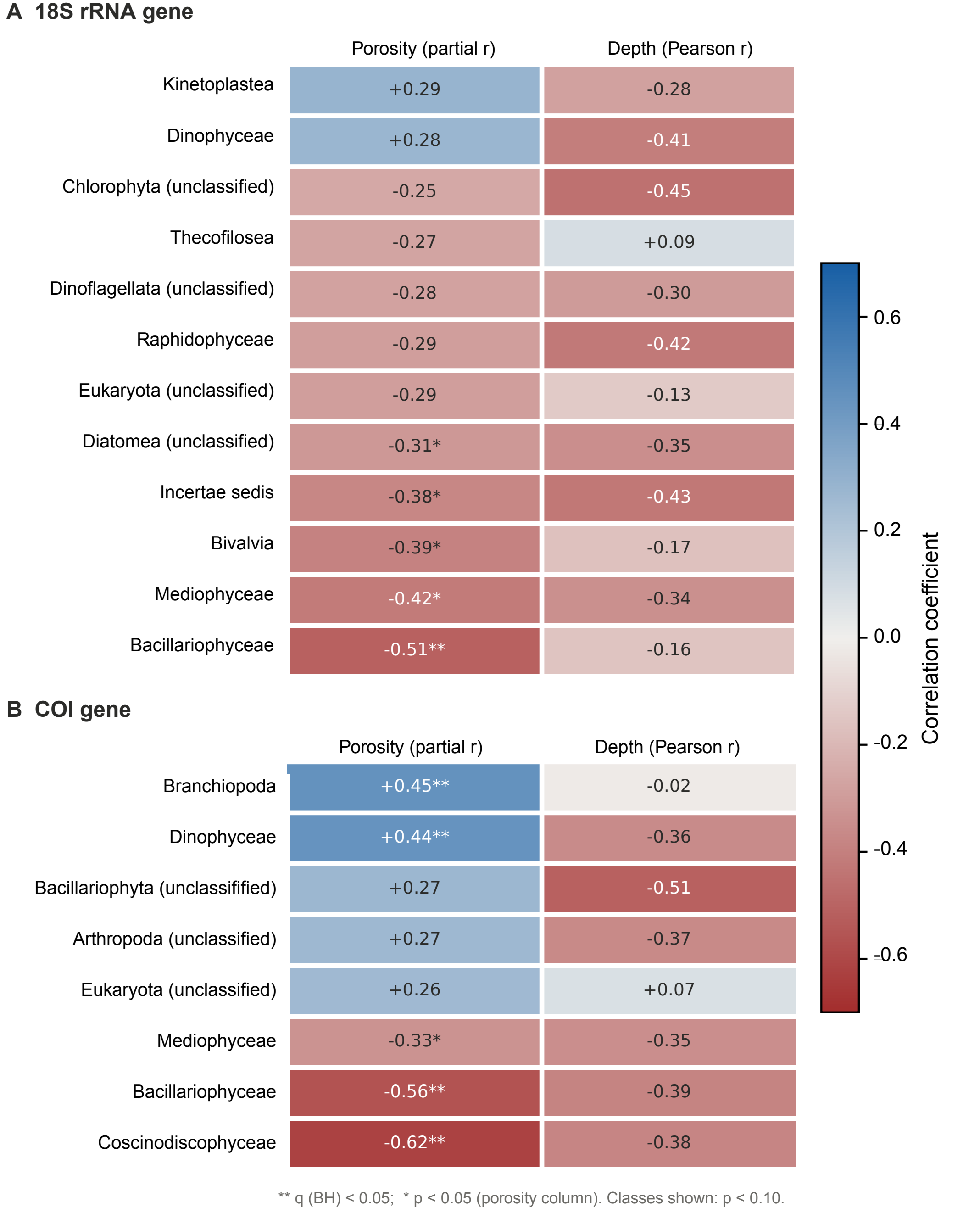


**Fig. S7.** Comparison of class-level correlations with depth-adjusted porosity and with burial depth for (A) the 18S rRNA gene (n = 44) and (B) the COI gene (n = 46) datasets. The left column shows partial correlation coefficients between porosity residuals and arcsine square-root transformed relative abundance, controlling for burial depth and log(burial depth + 1), as in Fig. S5. The right column shows Pearson correlation coefficients between relative abundance and burial depth. Classes shown are those with $p < 0.10$ for porosity. Colours indicate the correlation coefficient. In the porosity column, ** denotes Benjamini–Hochberg $q < 0.05$ and * denotes $p < 0.05$.
